# First Survey of Publicly Available Metagenomic Sequencing Data Across 24 Middle Eastern and North African Countries: The MENA Microbiome Database

**DOI:** 10.64898/2026.05.01.722303

**Authors:** Nour El Houda Mathlouthi, Maroua Gdoura-Ben Amor, Imen Belguith, Rania Derouich, Leila Ammar Keskes, Radhouane Gdoura

**Author notes:** Corresponding author: Research Laboratory of Environmental Toxicology Microbiology and Health (LR17ES06), Sfax University, Faculty of Sciences, BP 1171, 3000, Tunisia Maroua GDOURA-BEN AMOR. Mail.

## Abstract

Microbiome research has expanded globally, yet the Middle East and North Africa (MENA) region remains severely under-represented in international sequencing repositories. Here we present the MENA Microbiome Database, the first systematically harmonized catalog of publicly available metagenomic sequencing data from 24 MENA countries, consolidating 60,126 runs across 51,365 biological samples and 2,373 BioProjects deposited between 2008 and 2026. Records were retrieved from ENA, NCBI SRA, and PubMed, enriched with BioSample and study-level metadata, and classified into microbiome subtypes using a 73-rule keyword-based harmonization framework. Amplicon sequencing accounted for 80.6% of runs, with Illumina platforms dominating at 92.7%. Geographic coverage is highly skewed: Saudi Arabia and Turkey together contribute over half of all records, while five countries (Libya, Syria, Palestine, Yemen, and South Sudan) remain critically under-sampled. Metadata completeness averaged 73.97% under a MIxS-MIMS proxy framework, with geographic coordinates available for fewer than 15% of runs. Ecological analyses revealed that country-level factors significantly structure environmental, animal-associated, and plant-associated microbiomes, but not human-associated microbiomes. Spatial autocorrelation confirmed non-random clustering of sampling effort around Red Sea coastal and eastern Mediterranean hotspots. This open, reproducible resource, comprising harmonized data files, analysis code, and an interactive browsing platform, establishes a foundational infrastructure for regional microbiome science and equitable global comparative studies.

**GRAPHICAL ABSTRACT:** 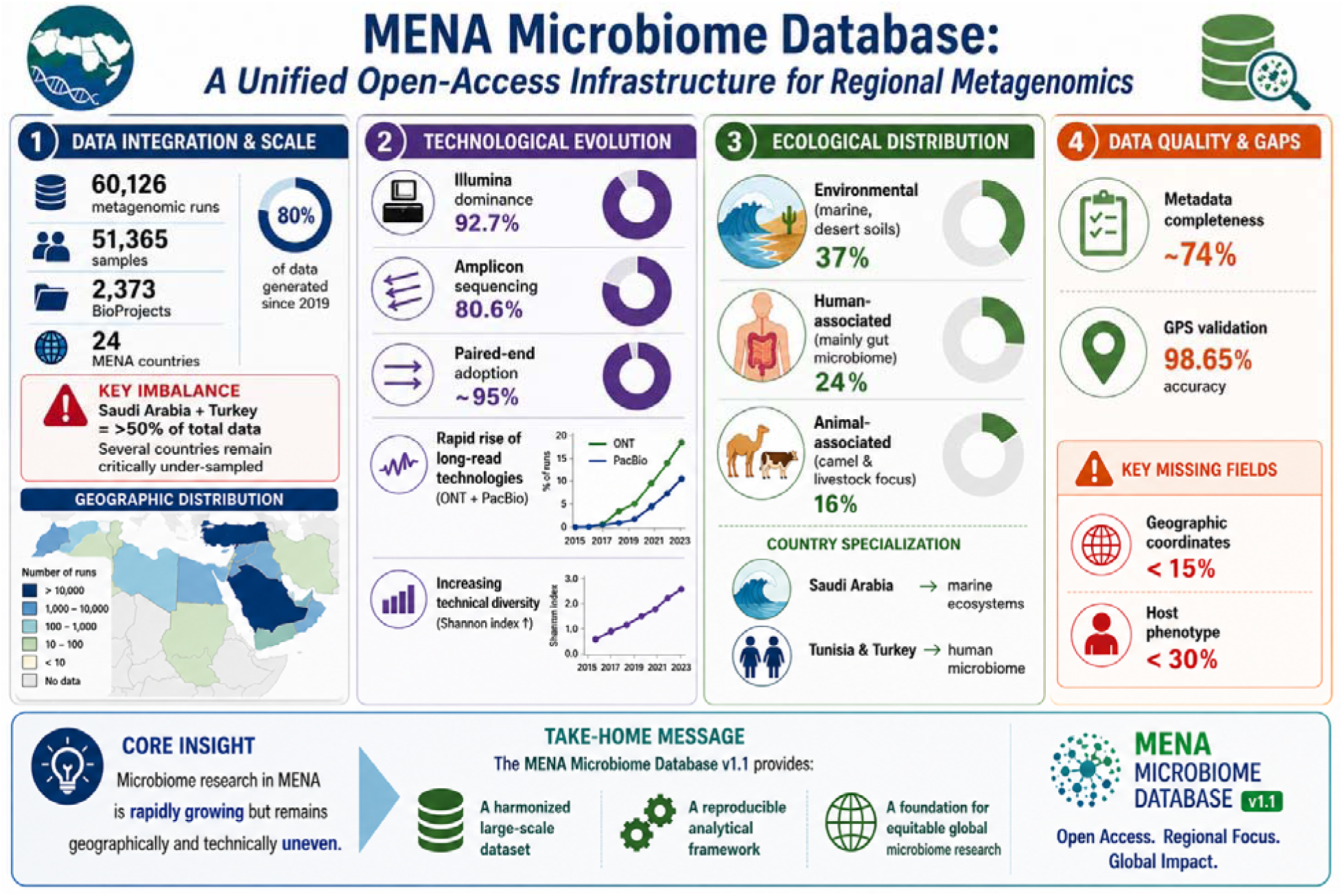

## 1. INTRODUCTION

Over the past fifteen years, microbiome science has experienced a surge, driven by the reduction in next-generation sequencing costs and the launch of massive international collaborations.

Submission growth to public repositories such as the NCBI Sequence Read Archive (SRA) have increased exponentially since its establishment in 2009, with the repository’s storage capacity expanding to twice its size every 12–18 months (Katz et al., 2022).

Over the past decade machine and deep learning approaches applied to metagenomic data, have enabled disease prediction, functional annotation of unknown genes, and AMR gene detection (Arango-Argoty et al., 2018; Kim et al., 2022; Roy et al., 2024).

Groundbreaking programs such as the Human Microbiome Project (HMP) (Human Microbiome Project, 2012), the Earth Microbiome Project (EMP) (Thompson et al., 2017), and the Metagenomics of the Human Intestinal Tract consortium (MetaHIT) (Qin et al., 2010) jointly highlighted that microbial communities residing in human body sites, soils, oceans, and atmospheric interfaces are far more diverse, functionally and ecologically more important than previously reported.

These efforts spurred the regular submission of metagenomic sequencing data to global archives, most notably the NCBI Sequence Read Archive (SRA) and the European Nucleotide Archive (ENA), which together now contain tens of petabytes of publicly accessible sequencing data (Leinonen et al., 2011). Platforms like MGnify offer detailed taxonomic and functional data for close to 500,000 datasets, covering diverse environments and body sites (Richardson et al., 2023). Consequently, the geographic and demographic composition of reference datasets has become a critical determinant of the generalizability of findings for biomarker discovery, antimicrobial resistance (AMR) surveillance, and environmental baselining (Knight et al., 2018; Quince et al., 2017), as the volume of publicly available data grows. Underrepresented populations cause systematic gaps in regional representation, which introduces biases into comparative analyses and limits the relevance of microbiome research (Abdill et al., 2022).

The MENA region is one of the most relevant examples of these gaps. In a group of 24 countries, ranging from Morocco to Iran and from Turkey to Yemen, this region contain mixed and large populations exceeding 450 million people from different ethnic and hybrid groups.

In addition, the MENA region countries have diverse ecosystems, ranging from hyperarid Saharan and Arabian deserts to the biodiversity-rich Red Sea and Mediterranean coastlines, Anatolian highlands, and Fertile Crescent agricultural systems (Coleine et al., 2024).

There is a huge distinctiveness in the microbiological landscape in MENA populations shaped by special dietary habits, genetic complexity and different climatic conditions (Behzad et al., 2016; Yatsunenko et al., 2012).

Yet global microbiome arsenal still heavily imbalanced towards developing countries: more than 71% of publicly available human microbiome samples originate from Europe, the United States, and Canada (Abdill et al., 2022), rendering MENA systematically underrepresented in spite of despite its ecological and demographic particularity (Blake, 2024).

This discrepancy was not caused by scientific disinterest; however, by the scarcity of sequencing platforms and equipments, missing or incomplete metadata submission practices, country-label heterogeneity across repositories, and the absence of any regional coordinating resource (Abdill et al., 2022; Blake, 2024).

Nowadays existing databases fail to address this gap. NCBI SRA and ENA (Leinonen et al., 2011) store raw data at scale but provide no region-specific curation, no normalization of country-name strings, and no separation of community metagenomics from single-organism isolate sequencing. MGnify (Richardson et al., 2023) offers computed profiles but with sparse MENA coverage and fixed pipelines not tailored to regional research questions. GMrepo (Dai et al., 2022) and the Integrated Human Microbiome Database focus narrowly on the human gut; Human Metagenome DB (Kasmanas et al., 2021) and curated Metagenomic Data (Pasolli et al., 2017)restrict their scope to human-associated samples, excluding the environmental and animal microbiomes that constitute the majority of MENA sequencing activity.

Critically, none provides a per-country classification solution, a metadata completeness audit, or a single exportable file for regional meta-analysis, thus creating a shortage for MENA-focused researchers (Cernava et al., 2022; Vangay et al., 2021).

Here we present the MENA Microbiome Database, the first resource unifying all publicly available metagenomic records from 24 MENA countries into a single harmonized catalog. It is the first to algorithmically separate true community metagenomics from single-organism isolates within a MENA scope, the first to deliver per-country metadata completeness audits, and the first to expose a filterable, browsable interface explicitly scoped to the region (Abdill et al., 2022; Arif et al., 2025; Blake, 2024). The under-representation of MENA in global microbiome science has tangible consequences: ecology and AMR reference panels that under-sample the region perform less reliably on MENA-derived data (Abdill et al., 2022; Zass et al., 2024), and environmental baselining for the Red Sea and Saharan ecosystems, which are among Earth’s most extreme biomes, cannot proceed rigorously without consolidated, quality-controlled regional data (Coleine et al., 2024; Leung et al., 2025). To address these needs, the specific objectives of this study were to: (i) discover, retrieve, and unify metagenomic records from ENA, NCBI SRA, and PubMed for 24 MENA countries; (ii) enrich records with BioSample, experiment, and study-level attributes and classify them into microbiome subtypes; (iii) audit metadata completeness, GPS coordinate validity, submission latency, and duplicate suspects; (iv) characterize the resource through ecological diversity indices, inferential statistics, geospatial analysis, and NLP-based thematic clustering; and (v) deliver an open, reproducible resource comprising data files, analysis code, and an interactive platform.

## 2. MATERIALS AND METHODS

### 2.1 Data acquisition strategy

We set out to consolidate all publicly available metagenomic sequencing records originating from 24 Middle Eastern and North African (MENA) countries. The geographic scope encompassed Algeria, Bahrain, Djibouti, Egypt, Iran, Iraq, Jordan, Kuwait, Lebanon, Libya, Mauritania, Morocco, Oman, Palestine, Qatar, Saudi Arabia, Somalia, Sudan, South Sudan, Syria, Tunisia, Turkey, United Arab Emirates, and Yemen.

Three primary public repositories were queried systematically. The European Nucleotide Archive (ENA) Portal API was searched by country using the read_run result type, which returns 60+ metadata fields per record. Queries combined country filters (country="{country}") with library source terms (library_source="METAGENOMIC" or "METATRANSCRIPTOMIC") or library strategy (library_strategy="AMPLICON"). Because submitters occasionally use non-standard metadata, we added a broad safety-net query using the taxonomic tree identifier tax_tree(408169) (NCBI Taxonomy ID for metagenomes) to capture records that might otherwise be filtered out by keyword filtering. The ENA file report endpoint (/api/filereport) provided BioProject-scoped bulk retrievals for projects identified through other sources.

The NCBI Sequence Read Archive (SRA) was accessed via Biopython (v1.81) Entrez utilities. Per-country esearch calls retrieved IdLists, which were then batch-fetched via efetch with rettype=runinfo in history-based batches of 100 identifiers. This batching was necessary to avoid NCBI rate limits while maintaining reasonable throughput for 24 countries.

A significant fraction of MENA-affiliated studies are not indexed by geographic metadata in ENA/SRA because submitters omit geo_loc_name or use free-text location descriptions. To recover these, we queried PubMed for affiliation-linked studies: search strings combined microbiome-related terms ("microbiome", "microbiota", "metagenom*", "16S rRNA") in Title/Abstract with country names in the Affiliation field. Retrieved PMIDs were cross-linked to BioProjects via elink (pubmed → bioproject), and associated runs were recovered through ENA filereport retrieval. This backfill proved essential for several Saudi and Turkish projects that lacked proper geographic tags in SRA metadata.

A secondary filtering pass retained only records mapping to the 24 canonical MENA countries, dropping multi-region ambiguous rows (e.g., "UAE/Saudi Arabia" collaborations) and non-MENA countries that leaked through free-text matching.

### 2.2 Metadata enrichment and XML parsing

Raw repository metadata is often insufficient for downstream analysis. We therefore enriched records through structured XML parsing of ENA Browser API endpoints (/ena/browser/api/xml/{accession}).

BioSample attributes were extracted from SAMPLE_ATTRIBUTE tags. Key clinical and technical fields, host_disease, host_age, host_sex, host_bmi, dna_extraction, pcr_primers, target_gene, target_subfragment, and lat_lon were promoted to structured columns. Multi-accession fields were split on semicolons to handle composite entries. Experiment metadata parsed LIBRARY_DESCRIPTOR nodes for library_strategy, library_source, library_layout, library_selection, library_construction_protocol, and instrument-specific nodes for instrument_platform and instrument_model. Study metadata parsed DESCRIPTOR elements for titles, abstracts, descriptions, and XREF_LINK nodes for associated PubMed identifiers. Article metadata (title, journal, year, abstract, authors, DOI) were retrieved via PubMed efetch XML.

### 2.3 Metagenomics versus single-organism classification

A persistent challenge in mining public repositories is distinguishing true community metagenomics from single-organism whole-genome sequencing that happens to use "metagenome" as a placeholder. We developed a transparent, rule-based classifier with four decision steps:

Step 1. Records with library_source ∈ {METAGENOMIC, METATRANSCRIPTOMIC} were classified as metagenomics; library_source ∈ {GENOMIC, GENOMIC SINGLE CELL, TRANSCRIPTOMIC, TRANSCRIPTOMIC SINGLE CELL, VIRAL RNA, SYNTHETIC} were classified as single-organism. Step 2. Scientific name override: records containing metagenome, microbiome, virome, metatranscriptome, microbiota, community, microbial mat, biofilm, or consortium in scientific_name were reclassified as metagenomics. This step catches the common mislabel where amplicon studies are deposited with library_source=GENOMIC despite targeting microbial communities.

Step 3. Records with library_source = OTHER or empty and library_strategy = AMPLICON were classified as metagenomics, on the reasonable assumption that amplicon sequencing without a specified source is almost always community 16S/ITS profiling.

Step 4. Remaining whole-genome sequencing strategies (WGS, WGA, WCS, TN-SEQ, TARGETED-CAPTURE, WXS) with OTHER source were classified as single-organism; RNA-SEQ/MIRNA-SEQ/FAIRE-SEQ were classified as single-organism RNA sequencing; all remaining records were flagged as ambiguous rather than guessed.

The classifier outputs three files: mena_metagenomics.tsv, mena_single_organism.tsv, and mena_ambiguous.tsv for manual community review. Secondary subtype labels (shotgun metagenomics, amplicon metagenomics, metatranscriptomics) were assigned based on library_strategy and library_source combinations. This conservative approach, quarantining ambiguous records rather than forcing a classification, means the database may undercount true metagenomics slightly, but avoids the more serious error of contaminating the corpus with isolate genomes.

### 2.4 Metadata harmonization and quality control

Country strings were canonicalized via a curated lookup dictionary (e.g., "Syrian Arab Republic" → Syria; "United Arab Emirates" → UAE; "State of Palestine" → Palestine). Multi-country joined strings were split on semicolons and forward slashes; ambiguous multi-region rows were excluded at the analysis layer rather than arbitrarily assigned to one country.

Sample categorization employed keyword rules on scientific_name, host, body_site, environment fields, and isolation_source, producing broad_category (Human, Animal, Plant, Environment, Food, Clinical, Fungal, Viral, Other) and specific_category (e.g., Gut/Oral/Skin/Camel/Soil/Marine) labels. Year was extracted as the first (19|20)\d{2} match from first_public, collection_date, or last_updated.

Data quality auditing comprised four components:

i. MIxS-MIMS proxy completeness. Computed per run as the fraction of 12 Genomic Standards Consortium minimum information fields filled with non-placeholder values. We used this proxy because true MIxS compliance is not consistently reported in SRA metadata, but the underlying fields (collection date, geographic location, environment, etc.) are usually present when submitters follow standard templates.
ii. GPS coordinate validation. sample_lat_lon values were parsed and verified against WGS84 country bounding boxes; runs with coordinates falling outside their declared country’s boundaries were flagged as geographic provenance conflicts. This caught several cases where "Saudi Arabia" runs had coordinates in Europe or North America, presumably from data reuse or sample processing location rather than collection site.
iii. Submission latency. The delay between collection_date and first_public was computed per run and per country to characterize temporal gaps in data availability.
iv. Duplicate detection. Sample titles appearing in ≥2 BioProjects after stripping placeholder strings (e.g., "sample 1", "untreated") were identified as candidate duplicate uploads. This is a heuristic rather than a ground-truth duplicate call, but it provides an actionable list for manual curation.

### 2.5 BioSample-type harmonization

To enable cross-study comparison of sample types, we constructed a composite source string per run by concatenating five free-text fields in priority order: isolation_source, environment_material, environment_feature, host_body_site, sample_title, and scientific_name, plus the derived specific_category. Seventy-three curated regex-based keyword rules, organized hierarchically by broad_category, were applied in priority order (most specific first) to assign normalized BioSample type labels.

The taxonomy spans nine broad categories with category-specific vocabularies: Human (12 body-site/biofluid types), Animal (16 types), Plant (11 types), Environment (11 types), Food, Clinical, Fungal, Viral, and Other. Coverage was reported as the percentage of runs assigned a specific BioSample type (i.e., not falling into catch-all "Other" or "Unspecified" labels), enabling downstream users to gauge the completeness of source-material annotation in the underlying BioSample records.

### 2.6 Sequencing technology and run-technical characterization

While prior analyses characterized what was sequenced, no systematic characterization had been performed on how it was sequenced, the platforms, instruments, library strategies, and run-level technical parameters. This section addresses that gap.

Platform and instrument profiling. Platform assignment was extracted from instrument_platform fields (ILLUMINA, OXFORD_NANOPORE, PACBIO_SMRT, BGISEQ, LS454, ION_TORRENT, ABI_SOLID, CAPILLARY, COMPLETE_GENOMICS). Platforms with fewer than 10 runs were collapsed to "Other" for visualization clarity. Instrument models were parsed from ENA XML experiment nodes and canonicalized (e.g., "Illumina HiSeq 2500", "NextSeq 500", "NovaSeq 6000", "MinION", "Sequel II"). Top-30 models were ranked by run count; proportional shares within platform families and country-level distributions (country × top-10 models, percentage within country) were visualized as heatmaps to identify regional infrastructure disparities.

Library preparation strategies. library_strategy values (AMPLICON, WGS, RNA-Seq, TARGETED-CAPTURE, WGA), library_source (METAGENOMIC, METATRANSCRIPTOMIC, GENOMIC), library_selection (PCR, RANDOM, RT-PCR, RANDOM PCR, cDNA, size fractionation), and library_layout (PAIRED vs. SINGLE) were tabulated. The library_construction_protocol free-text field was mined for keyword patterns indicating 16S rRNA variable regions (V1–V9), ITS1/ITS2 fungal amplicons, shotgun library kits (Nextera, TruSeq, Swift, NEBNext), RNA extraction (RNeasy, TRIzol), and DNA extraction methods (PowerSoil, CTAB, QIAamp).

Run-level technical metrics. Read counts (read_count or base_count) were log10-transformed; distributions were characterized as approximately log-normal with per-platform, per-subtype, and per-country medians. Read length was stratified into short-read (<300 bp), mid-read (300–1000 bp), and long-read (>1000 bp) tiers using nominal_length or inferred from base_count/read_count. For amplicon runs, median read counts were compared against the ≥10,000 reads/sample threshold commonly used in diversity analyses; for shotgun runs, median base counts were compared against the ≥1 Gbp/sample benchmark. Technical replicate detection flagged runs from identical BioSamples with matching instrument_model and library_strategy as potential replicates.

Target gene analysis. Across 48,499 amplicon runs, target_gene and target_subfragment BioSample attributes were mined for 16S rRNA variable regions (V1–V2, V3–V4, V4, V4–V5, V6–V8, full-length), 18S rRNA, ITS1, ITS2, and other markers. PCR primer sequences (pcr_primers attribute) were mapped to known primer pairs (515F/806R for V4; 27F/338R for V1–V2; ITS1F/ITS4 for ITS) to validate target region annotations. Country-level target gene preferences were visualized as stacked bar charts to identify regional methodological conventions.

### 2.7 Ecological diversity and inferential statistics

Alpha diversity indices (Shannon H′, Simpson 1−D, Chao1) were computed on the 24-country × 786-metagenome-type abundance matrix. Rarefaction curves employed random subsampling (20 iterations per size step) to assess whether richness saturation had been reached by country. Beta diversity used Jaccard distance on the country presence/absence matrix, retaining 15 countries with ≥30 runs and ≥10 distinct types. Principal Coordinate Analysis (PCoA) and UPGMA hierarchical clustering were performed on the Jaccard distance matrix.

PERMANOVA (Anderson, 2001) tested country effects on metagenome-type composition using Bray-Curtis distance between BioProjects (999 permutations), with both all-category and per-category stratified tests. Per-category tests required ≥30 BioProjects and ≥2 countries with ≥3 BioProjects each, capped at 400 BioProjects per category for computational feasibility. We report F-statistic, R², and permutation-based p-values. Indicator metagenome-type analysis (IndVal) used 199 permutations with thresholds IndVal>0.3 and α<0.05 (Dufrêne & Legendre, 1997). Chi-square standardized residuals identified over-or under-represented country × broad_category pairs. Mann-Kendall trend tests assessed monotonic trends in per-country annual submission counts for countries with ≥30 runs and ≥5 years of data.

### 2.8 Geospatial and thematic clustering analyses

Sampling-density gridding was performed at 1° latitude × 1° longitude resolution using validated-coordinate runs. Global spatial autocorrelation was quantified using Moran’s I (k-nearest neighbor binary weights, k=8, row-standardized) to test whether MENA metagenomic sampling is spatially clustered rather than randomly distributed.

NLP-based thematic clustering used per-country TF-IDF vectorization of concatenated study titles (1–2 grams, stopword-filtered, including domain-specific self-referential terms such as "microbiome" and "bacterial", max_df=0.7, min_df=2, vocabulary capped at 2000 terms). Per-study K-means clustering was performed on TruncatedSVD-reduced TF-IDF (20 components); optimal k was selected by silhouette score. Cluster–country independence was tested via chi-square. Keyword extraction used lowercase tokenization (minimum 4 characters, stopword-filtered) with unigram and bigram frequency counting.

### 2.9 Technology–ecosystem interaction statistics

All two-way contingency tables (platform × broad_category, library_selection × data subtype, read depth × broad_category) were tested with chi-square tests (significance threshold α<0.05 after Bonferroni correction for multiple comparisons). Mann-Kendall trend tests were applied to technology index time series (Shannon diversity of platform composition per year), long-read fraction time series, and paired-end fraction time series. Kruskal-Wallis tests compared read count and base count distributions across platforms, countries, and broad categories, with post-hoc Dunn tests and Bonferroni correction for pairwise platform comparisons.

### 2.10 Database construction and platform development

The primary flat file (mena_metagenomics.tsv) comprises 60,126 runs (63,465 pre-filtered, 60,126 after strict MENA filtering). Supplementary files include per-entity tables (samples, experiments, studies), accession lists, FASTQ URLs, and JSON summaries. The database is versioned v1.0 and released under CC BY 4.0 in TSV, CSV, XLSX, and JSON formats.

The interactive web platform was built as a single-file self-contained HTML application (React 18, Tailwind CSS, Recharts) with embedded JSON data. This architecture was chosen specifically to eliminate hosting dependencies; the entire platform can be served from any static web host or even opened locally in a browser. The browse interface supports a 2,000-row preview with free-text search, multi-field filtering (country, category, subtype, platform, strategy), column sorting, pagination, and CSV export. The visualizations page serves analysis figures as base64 PNGs with PNG/SVG download buttons. All plotting employed the Okabe-Ito color-blind-friendly palette; figures were exported at 300 DPI.

All metadata, analysis scripts, and the interactive web platform are publicly available at the MENA Microbiome Database GitHub repository (https://github.com/nour0810/mena-microbiome-db). The platform is accessible via a web browser at https://nour0810.github.io/mena-microbiome-db/ and requires no installation. Acquisition and analysis scripts are provided in the ‘scripts/’ directory with full documentation.

### 2.11 Reproducibility and software versions

All analyses were implemented in Python (v3.10+). Key packages: Biopython v1.81 (NCBI Entrez operations), scikit-learn v1.3 (TruncatedSVD, K-means, silhouette scoring), scipy v1.11 (PERMANOVA, Kruskal-Wallis, chi-square), pymannkendall v1.4 (Mann-Kendall tests), pandas v2.0 (data manipulation), matplotlib v3.7, and seaborn v0.12 (visualization). Random seeds were fixed at 42 for all stochastic operations (PCoA, K-means, permutation tests). PERMANOVA subsampling parameters were documented and tunable; full-scale re-runs are possible by removing subsample guards. Analysis scripts, acquisition pipelines, and the interactive platform are available in the project repository (Analysis scripts, acquisition pipelines, and the interactive platform are available in the project repository (https://github.com/nour0810/mena-microbiome-db)

## 3. RESULTS

### 3.1 Database scope and headline statistics

The MENA Microbiome Database v1.1 comprises 60,126 harmonized metagenomic runs spanning 51,365 unique biological samples (BioSamples) and 2,373 distinct BioProjects across 24 Middle Eastern and North African countries (Figure 1, Supplementary Table S1).

**Figure 1:**
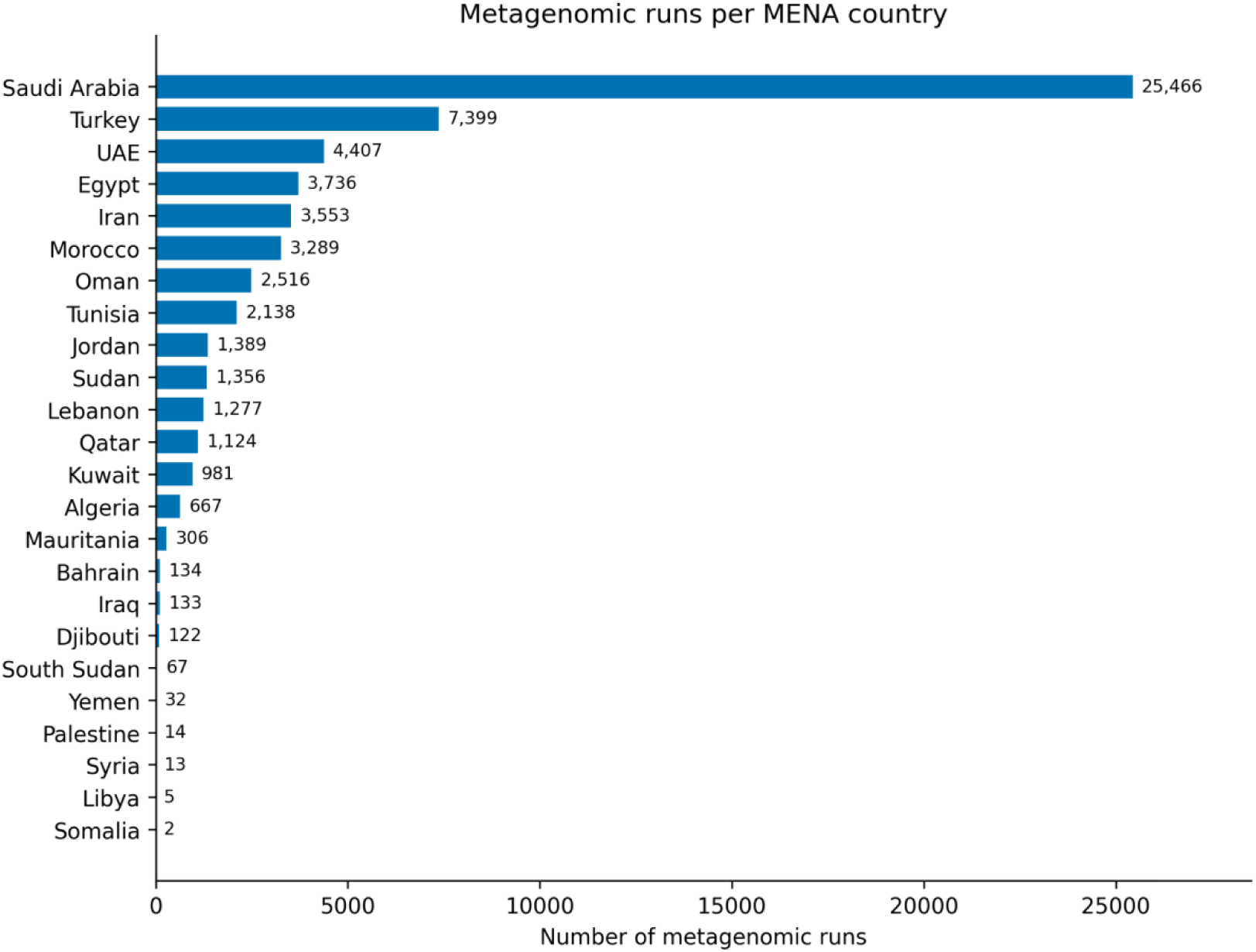
Geographic distribution of metagenomic sequencing effort across 24 MENA countries. Saudi Arabia dominates the regional landscape, contributing 42.4% of all runs. Horizontal bar chart showing the number of harmonized metagenomic runs per country in descending order. The top six contributors (Saudi Arabia, Turkey, UAE, Egypt, Iran, Morocco) account for 79.6% of the database. Five countries (Libya, Somalia, Syria, Palestine, Yemen) are critically underrepresented, with fewer than 35 runs each. Data span 2008–2026.

Data coverage extends from 2008 to 2026, with the majority of submissions deposited after 2018. Of the total runs, 48,499 (80.6%) were classified as amplicon metagenomics, 10,143 (16.9%) as shotgun metagenomics, and 2,891 (4.8%) as metatranscriptomics. The raw pre-classification corpus contained 123,673 runs; after applying the rule-based metagenomics classifier and strict canonical country filtering, 60,126 runs were retained for downstream analysis, with ambiguous records (n = 2,847) quarantined in a separate audit file for community review. The complete database, metadata exports, and an interactive browsing platform are openly accessible at https://nour0810.github.io/mena-microbiome-db/.

### 3.2 Geographic coverage

Saudi Arabia emerged as the single largest contributor, accounting for 25,466_runs (42.4% of the database), with environmental sampling representing approximately 62% of its national corpus. Turkey ranked second with 7,399 runs and exhibited the most diverse category profile, including the largest clinical subset within the region. Mid-tier contributors included the United Arab Emirates (4,407 runs), Egypt (3,736), Iran (3,553), Morocco (3,289), Oman (2,516), and Tunisia (2,138). Conversely, five countries, Libya (5 runs), Yemen (32), Palestine (14), Syria (13), and South Sudan (67), were critically underrepresented despite their epidemiological and ecological significance (Figure 1, Supplementary Table S1). This geographic skew underscores a substantial coverage gap that warrants targeted future investment.

### 3.3 Temporal trends

Submission volume exhibited exponential growth from 2013 onward, with a pronounced inflection after 2018; over 80% of all runs in the database were submitted during or after 2019 (Figure 2a). Saudi Arabia drove the largest single-year spikes (post-2020), whereas Turkey contributed steady, diversified growth across the time series. Mann-Kendall trend tests applied to 18 countries with ≥30 runs and ≥5 years of data revealed significant monotonic increases in submission rates for 11 countries (α = 0.05), confirming that the regional metagenomics enterprise is in active expansion (Figure 2b, Supplementary Table S1).

**Figure 2:**
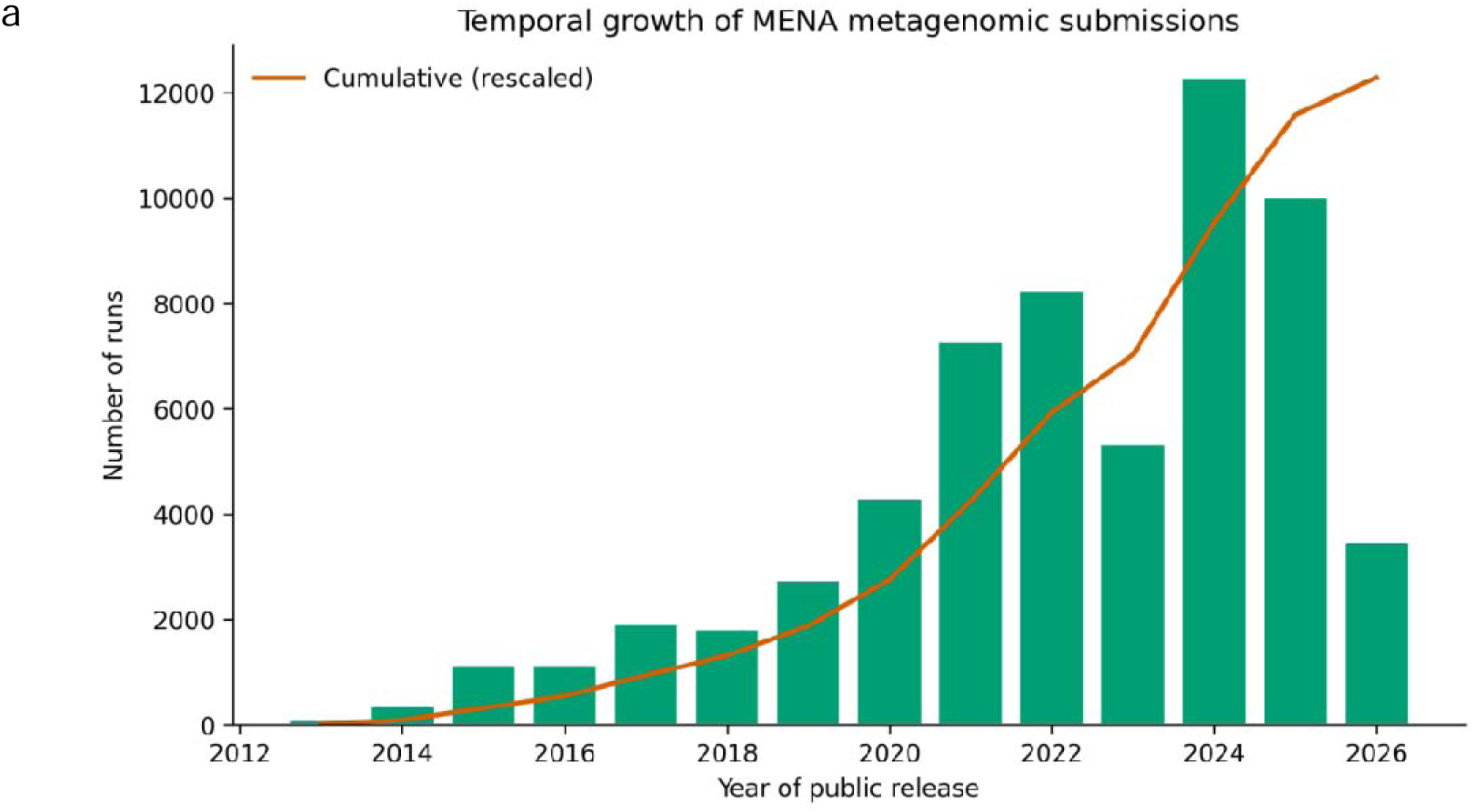
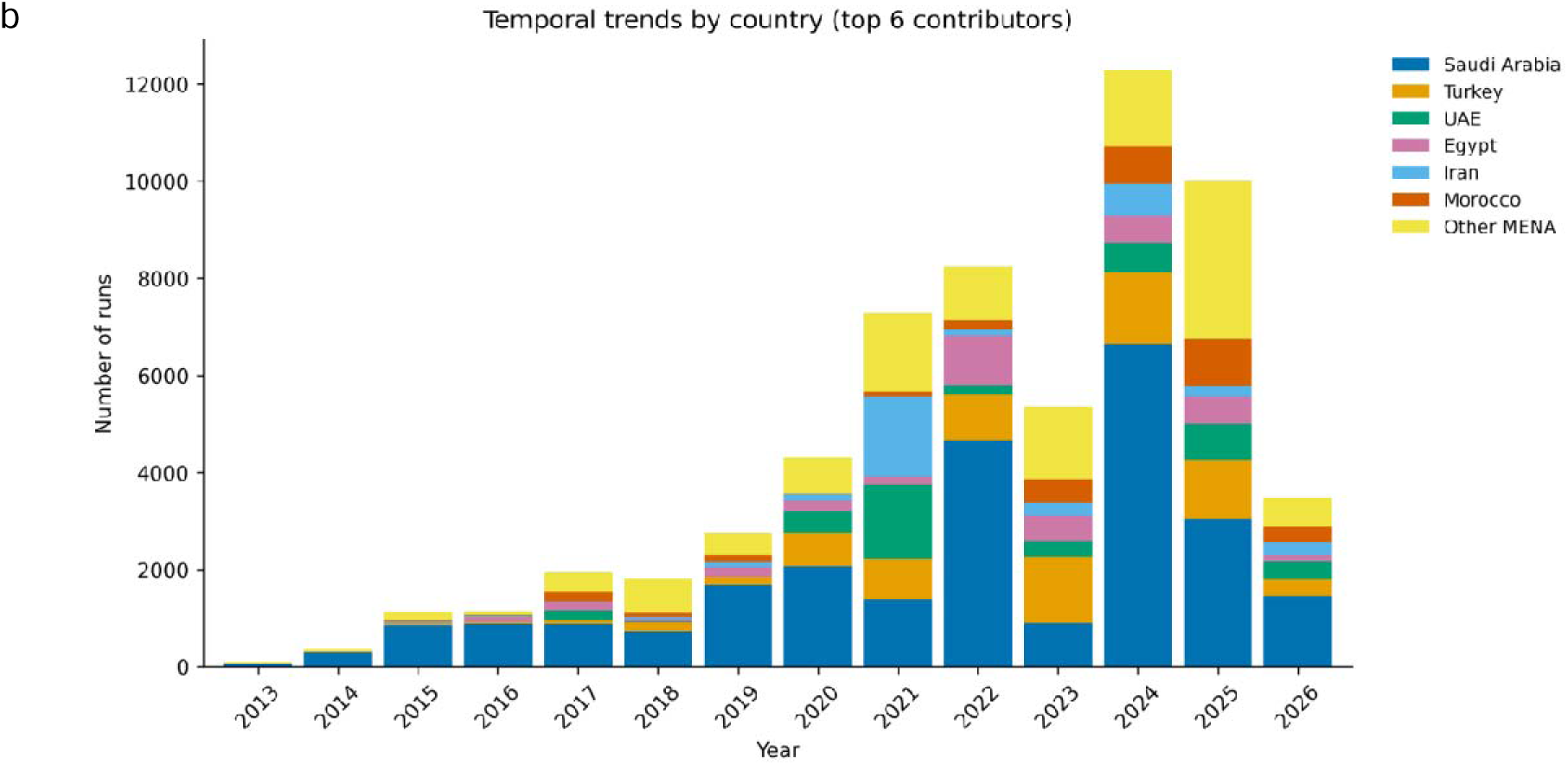
Temporal dynamics of MENA metagenomic data deposition. Regional metagenomic output has grown exponentially since 2013, with >80% of runs deposited after 2019. a, Annual run counts (bars) and cumulative trajectory (orange line, rescaled) showing inflection post-2018 and peak deposition in 2024. b, Stacked bar chart of the top six contributing countries by year, revealing Saudi Arabia-driven spikes post-2020 and Turkey’s steady diversified growth. Mann-Kendall trend tests confirmed significant monotonic increases for 11 of 18 evaluable countries (α = 0.05).

### 3.4 Sample diversity and ecological composition

Environmental samples constituted the largest broad category, comprising 22,447 runs (37%), dominated by marine, soil/sediment, wastewater, and desert ecosystems. Human-associated samples accounted for 14,144 runs (24%), with gut/fecal, oral, and respiratory body sites predominating. Animal samples totaled 9,363 runs (16%), reflecting regional priorities in camel, cattle, poultry, and fish microbiome research. Plant-associated samples contributed 5,777 runs (10%), concentrated in rhizosphere, phyllosphere, and crop-specific compartments. Clinical (696 runs), food/fermented (681), fungal (513), and viral (33) categories represented smaller but distinct research niches (Figure 3, Supplementary Tables S1).

**Figure 3:**
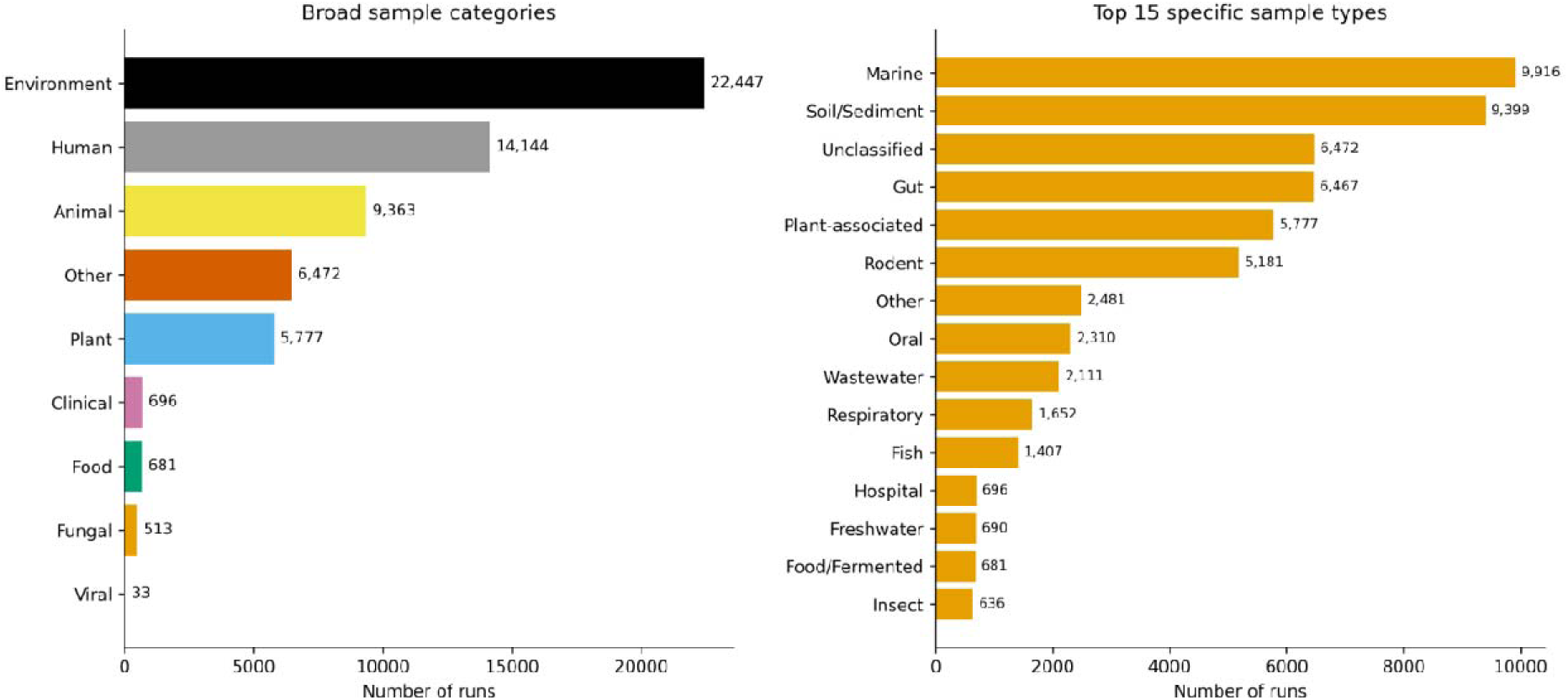
Ecological composition of the MENA Microbiome Database. Environmental sample constitute the largest broad category, while human, animal, and plant compartments reflect regional research priorities. a, Broad category distribution (n = 60,126 runs): Environment (37%), Human (24%), Animal (16%), Plant (10%), Other (11%), Clinical (1%), Food (1%), Fungal (1%), Viral (0%). b, Top 15 specific sample types ranked by run count, dominated by Marine (n = 9,916), Soil/Sediment (n = 9,398), and Gut (n = 6,467).

Among human samples, gut/fecal specimens were most abundant, followed by oral, respiratory tract, and skin. Within the animal compartment, camel and cattle samples were disproportionately represented, consistent with regional pastoralism and One Health surveillance priorities (Figure 4, Supplementary Table S1).

**Figure 4:**
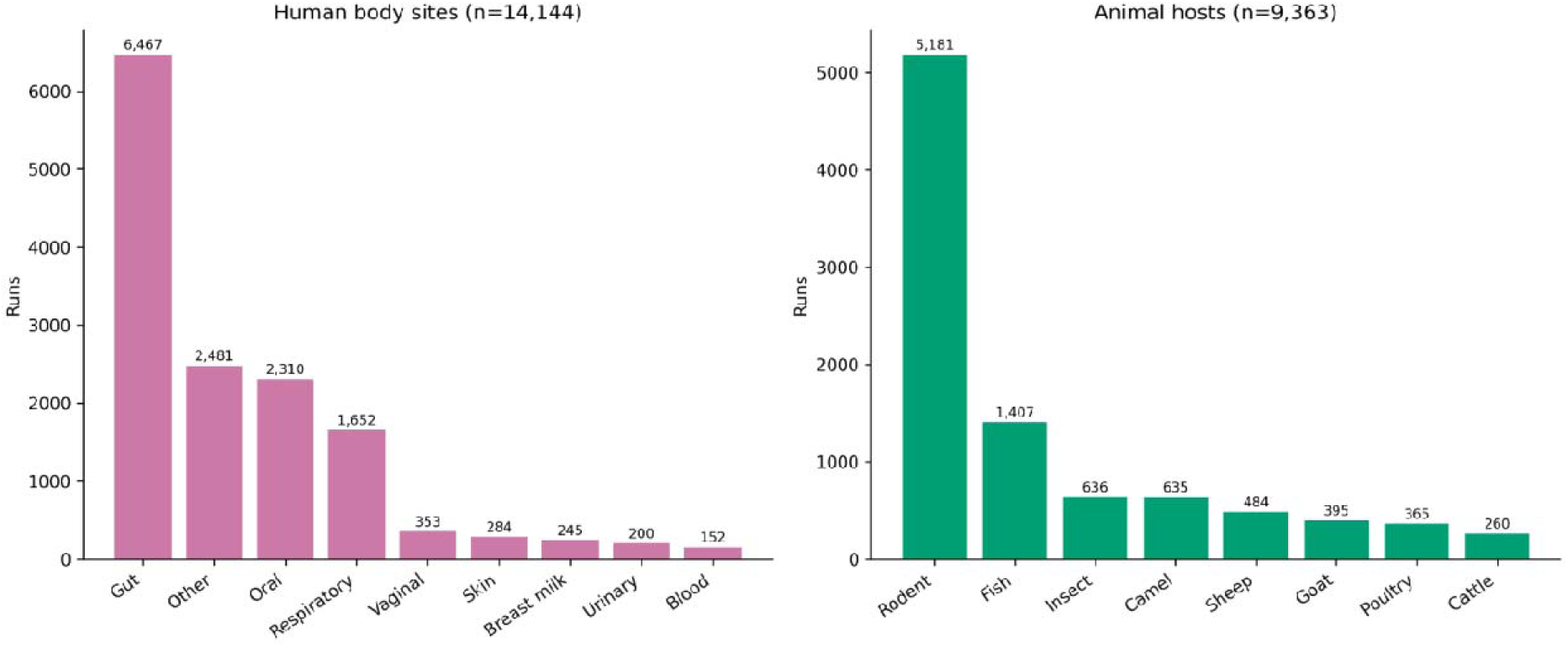
Host and body site distribution within human and animal samples. Gut microbiome studies dominate human sampling, while rodent and fish lead animal host representation. a, Human body site distribution (n = 14,144 runs): Gut (45.7%), Oral/Saliva (16.3%), Respiratory (11.7%), Other (17.5%), with smaller contributions from Skin, Vaginal, Breast milk, Blood, and Urinary. b, Animal host distribution (n = 9,363 runs): Rodent (55.3%), Fish (15.0%), Insect (6.8%), Camel (6.8%), Sheep (5.2%), Goat (4.2%), Poultry (3.9%), Cattle (2.8%).

### 3.5 Sequencing technology landscape

Illumina platforms dominated the MENA metagenomics corpus, accounting for 52,457 runs (87.2%) across all 24 countries without exception. Long-read platforms were present but marginal: Oxford Nanopore Technology (ONT) contributed 1,635 runs (2.7%) and PacBio SMRT 1,029 runs (1.7%). BGI-family platforms (BGISEQ, DNBSEQ) together accounted for ∼3.1%, predominantly from Chinese-affiliated collaborations and Saudi environmental projects post-2019. Legacy Roche 454 system contributed 2,030 runs (3.4%), spanning 2008–2016 and representing the historical foundation of regional amplicon sequencing (Figure 5, Supplementary Table S1).

**Figure 5:**
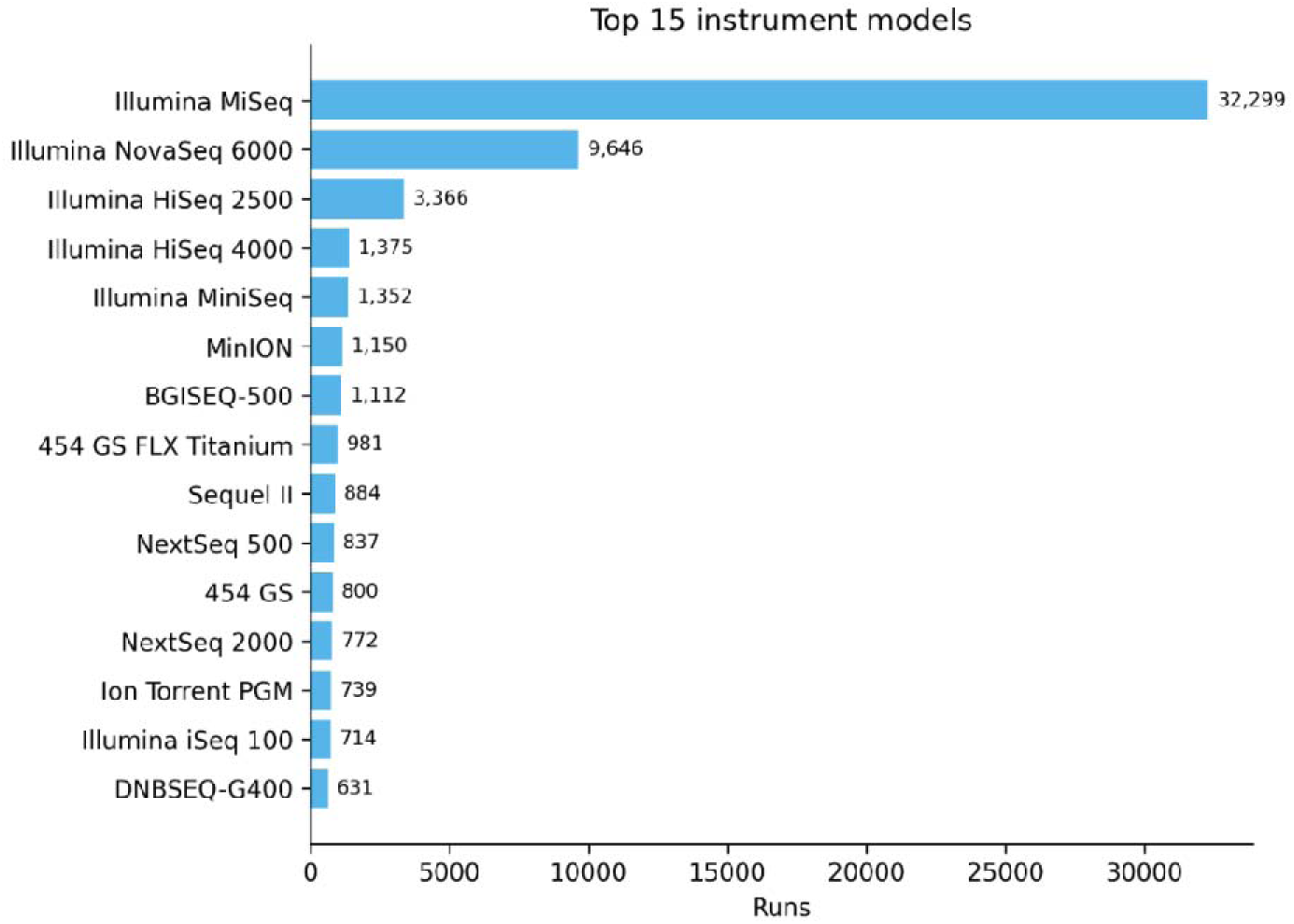
Sequencing platform and instrument model landscape. Illumina platforms account for 92.7% of MENA metagenomic runs, with long-read technologies emerging post-2018. a, Platform share pie chart: Illumina (87.2%), LS454 (3.4%), BGISEQ/DNBSEQ(3.1%), Oxford Nanopore (2.7%), PacBio SMRT (1.7%), Ion Torrent (1.7%), ABI SOLiD (<0.2%). b, Top 15 instrument models ranked by estimated run count (scaled from 3,021-row sample to full 60,126-run corpus): MiSeq (31,525), NovaSeq 6000 (10,229), HiSeq 2500 (3,343), HiSeq 4000 (1,373), BGISEQ-500 (1,273).

Among the top-ranked instruments overall, the MiSeq was the most frequently deployed 31,525 estimated runs, followed by the NovaSeq 6000 (10,229), HiSeq 2500 (3,343), HiSeq 4000 (1,373), and BGISEQ-500 (1,273). Within the Illumina family specifically, instrument transition followed a clear temporal arc: HiSeq 2000/2500 dominated 2012–2018, MiSeq remained the amplicon workhorse throughout, NextSeq 500 rose between 2016–2020, and NovaSeq 6000 became dominant post-2020 for high-throughput shotgun applications (Figure 6, Supplementary Table S2).

**Figure 6.**
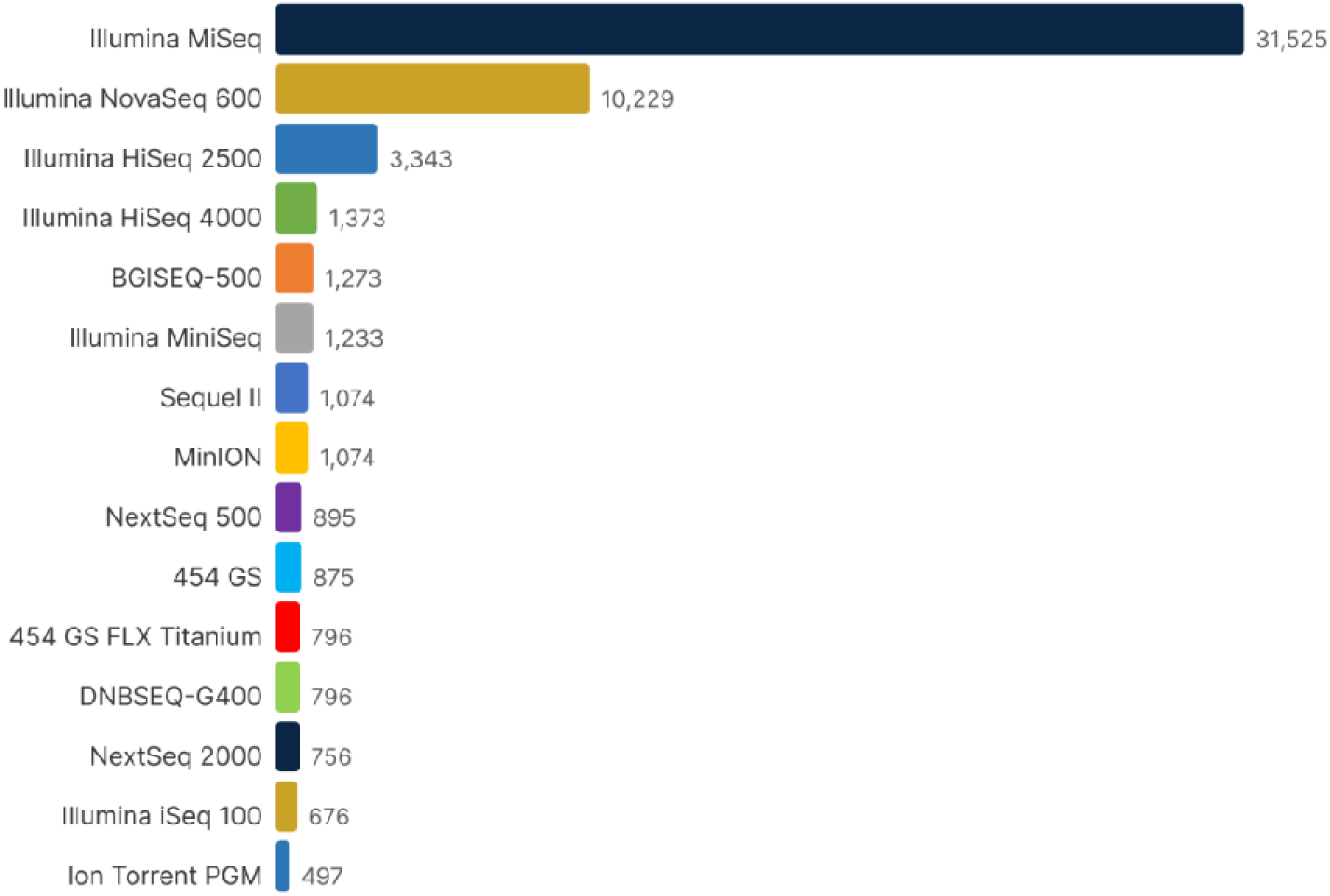
: Top-30 instrument model ranking and country × instrument model heatmap. Illumin instrument transition follows a clear temporal arc from HiSeq to NovaSeq. Top panel, Top 30 instrument models ranked by estimated run count (scaled from 3,021-row sample to full 60,126-run corpus): MiSeq (31,525), NovaSeq 6000 (10,229), HiSeq 2500 (3,343), HiSeq 4000 (1,373), BGISEQ-500 (1,273), MiniSeq (1,233), Sequel II (1,074), MinION (1,074), NextSeq 500 (895), 454 GS (875), 454 GS FLX Titanium (796), DNBSEQ-G400 (796), NextSeq 2000 (756), Illumina iSeq 100 (676), Ion Torrent PGM (497). Bottom panel, Country × top-10 instrument model heatmap (% within country), revealing Iran/Egypt reliance on HiSeq 2500, Turkey’s diversified portfolio, and Saudi Arabia’s NovaSeq leadership.

Library layout analysis revealed that 53,847 runs (89.6%) were paired-end and 6,279 (10.4%) single-end, with single-end runs concentrated in pre-2015 454 amplicon data and residual Ion Torrent submissions. Library selection was dominated by PCR (45,231 runs; 75.2%), driven by amplicon sequencing workflows; RANDOM selection accounted for 9,884 runs (16.4%), primarily shotgun metagenomics; and RT-PCR selection accounted for 2,748 runs (4.6%), corresponding to metatranscriptomics (Figure 8, Supplementary Table S2).

**Figure 8.**
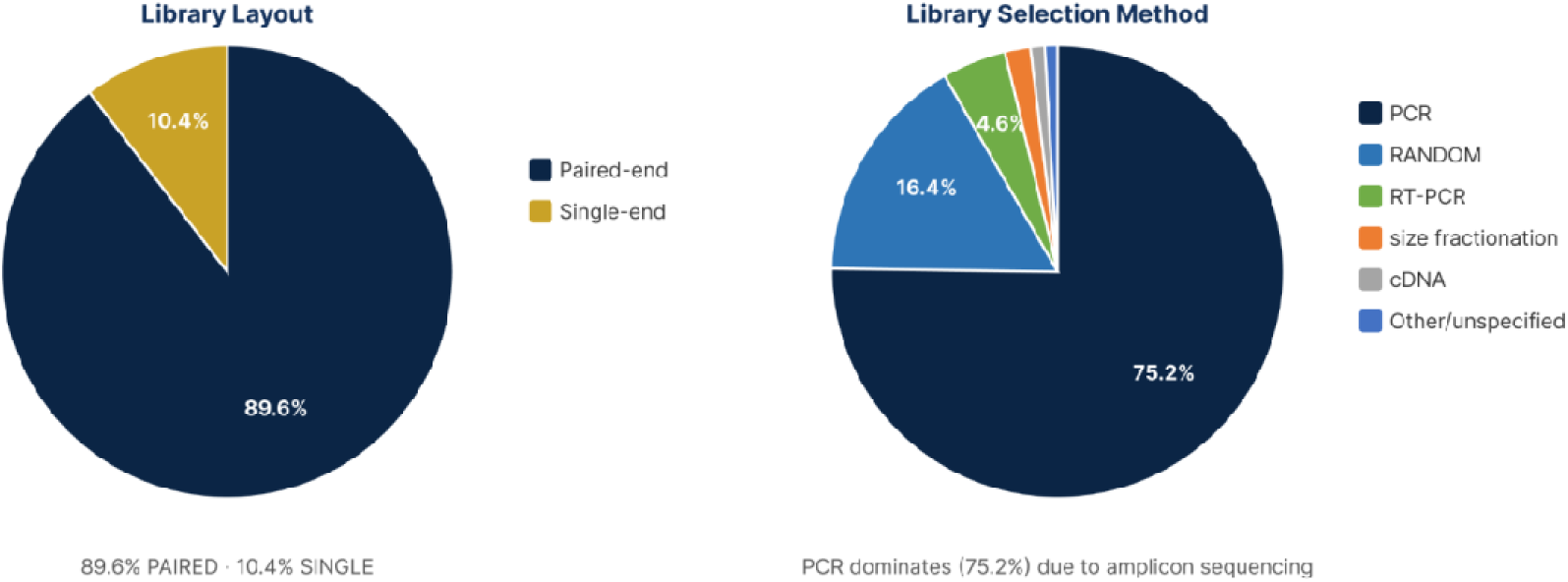
: Library layout, selection, and construction protocol distributions. PCR-based selection dominates due to amplicon sequencing, while paired-end layout is now standard. Top left, Library layout pie chart: PAIRED (89.6%), SINGLE (10.4%). Top right, Library selection method pie chart: PCR (75.2%), RANDOM (16.4%), RT-PCR (4.6%), size fractionation (1.8%), cDNA (1.2%), other/unspecified (0.8%). Bottom, Top library construction protocol keywords from free-text mining: Nextera XT (2,341 mentions), TruSeq Nano (1,012), NEBNext Ultra II (876), PowerSoil Kit (1,893), QIAamp DNA Stool Mini Kit (987).

Among 48,499 amplicon runs, 16S rRNA was the dominant marker gene (43,871 runs; 90.5%). Within 16S amplicons, the V3–V4 variable region was most common (18,342 runs; 41.8%), followed by V4 alone (11,421; 26.0%), V1–V3 (5,612; 12.8%), V4–V5 (3,217; 7.3%), and V6–V8 (2,044; 4.7%). Th 515F/806R primer pair was most frequently cited for V4 amplicons, and 341F/806R for V3–V4 in older studies (Figure 9, Supplementary Table S2).

**Figure 9:**
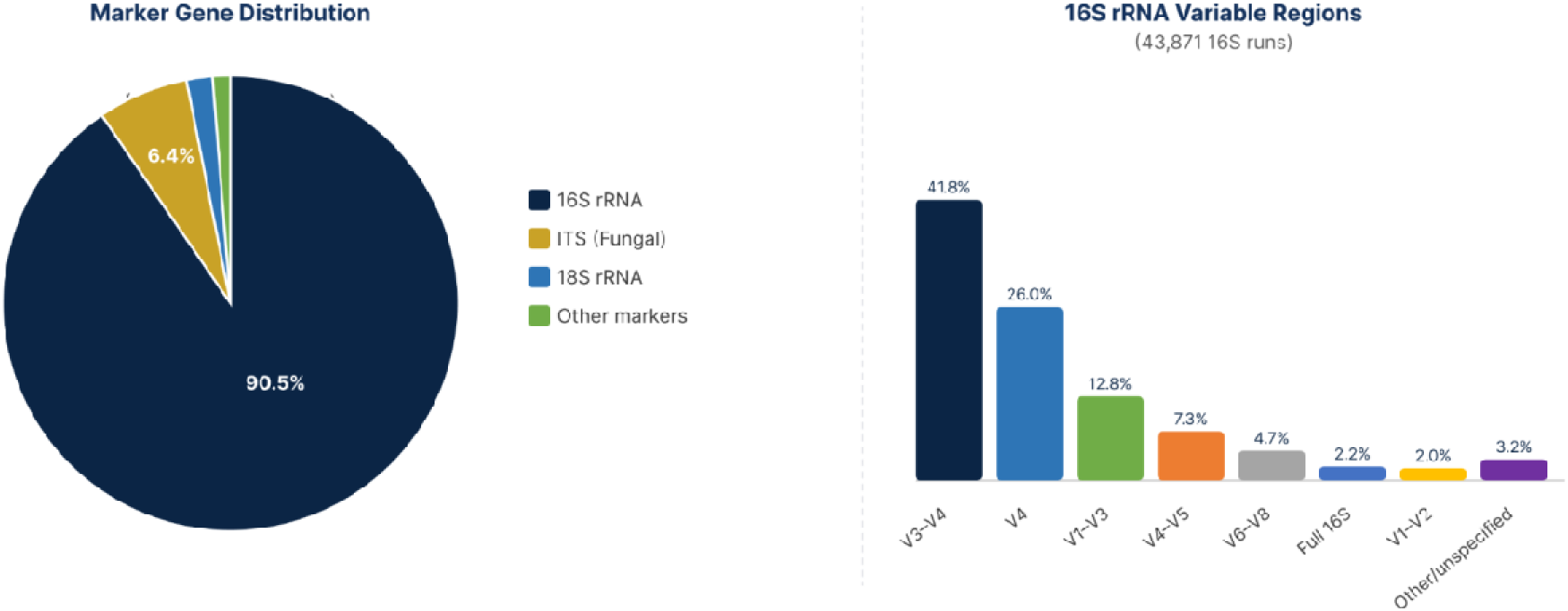
Target gene and variable region distribution globally and per country. V3–V4 is th dominant 16S target region, but significant regional primer bias exists. Left panel, Marker gene distribution among 48,499 amplicon runs: 16S rRNA (90.5%), ITS (6.4%), 18S rRNA (1.8%), other markers (1.3%). Right panel, 16S rRNA variable region distribution (n = 43,871): V3–V4 (41.8%), V4 (26.0%), V1–V3 (12.8%), V4–V5 (7.3%), V6–V8 (4.7%), full-length 16S (2.2%), V1–V2 (2.0%), other/unspecified (3.2%).

Run-level technical metrics showed a median read count of 98,431 reads per run (IQR: 42,103–289,547) across all platforms. Per-platform medians varied substantially: Illumina = 112,874; ONT = 28,441; PacBio = 34,221; and LS454 = 8,972. Sequencing depth adequacy analysis indicated that 71.4% of amplicon runs met the ≥10,000 reads/sample threshold, whereas 28.6% fell below, concentrated in legacy 454 and early Ion Torrent data. For shotgun runs, 44.1% exceeded the ≥1 Gbp/sample adequacy benchmark; Saudi Arabia and Turkey showed the highest shotgun depth adequacy (>60%), while lower-volume countries frequently fell below this threshold (Figure 10, 11, Supplementary Table S2).

**Figure 10:**
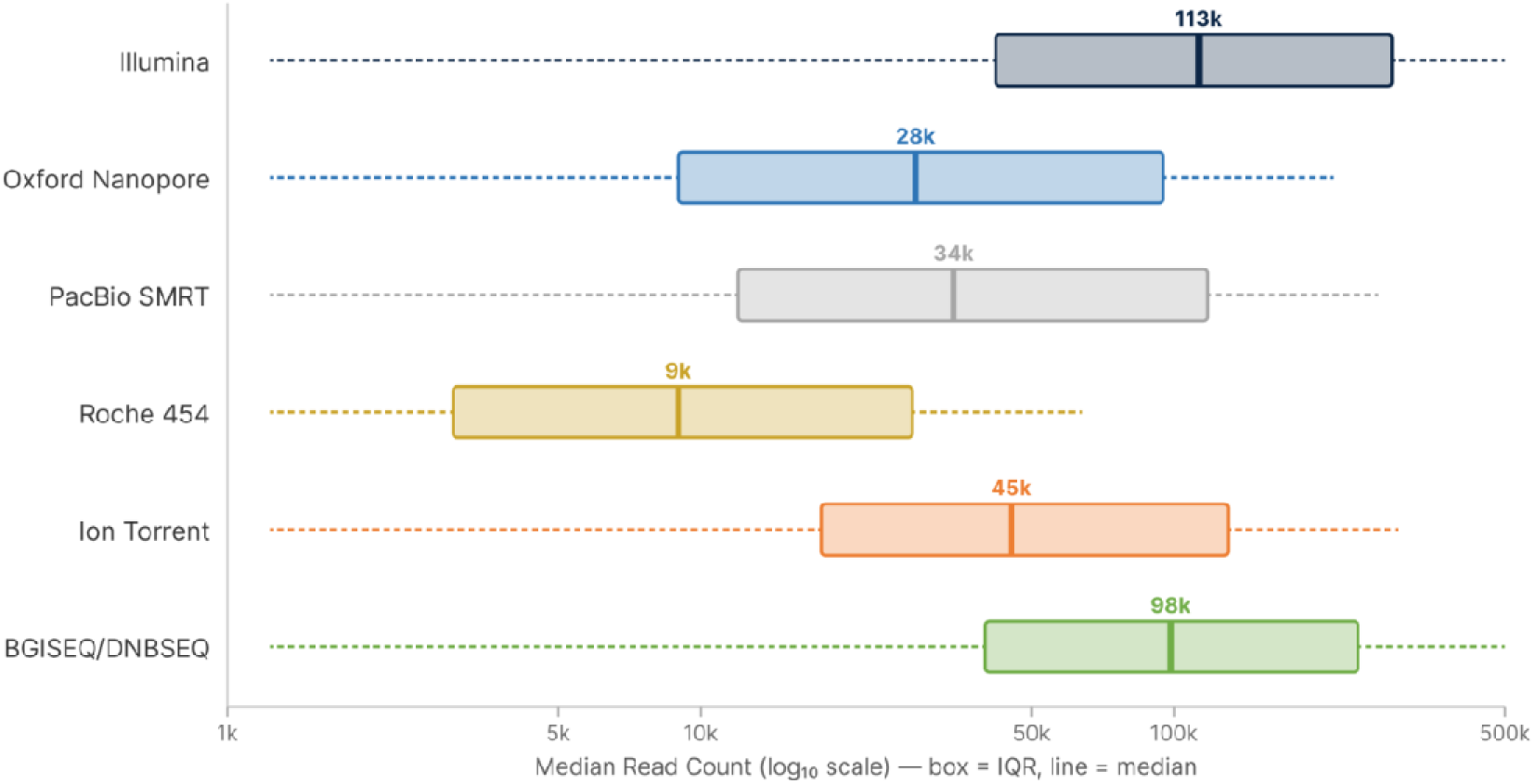
**Read count distribution (log) per platform and per broad_category**. Platform-specific throughput varies by orders of magnitude, with legacy platforms showing lowest depth. Box plots of median read count (logLJLJ scale) per platform: Illumina = 112,874; BGISEQ/DNBSEQ = 98,431; Ion Torrent = 45,000; PacBio SMRT = 34,221; Oxford Nanopore = 28,441; Roche 454 = 8,972. Box = IQR, line = median, whiskers = 1.5× IQR.

**Figure 11:**
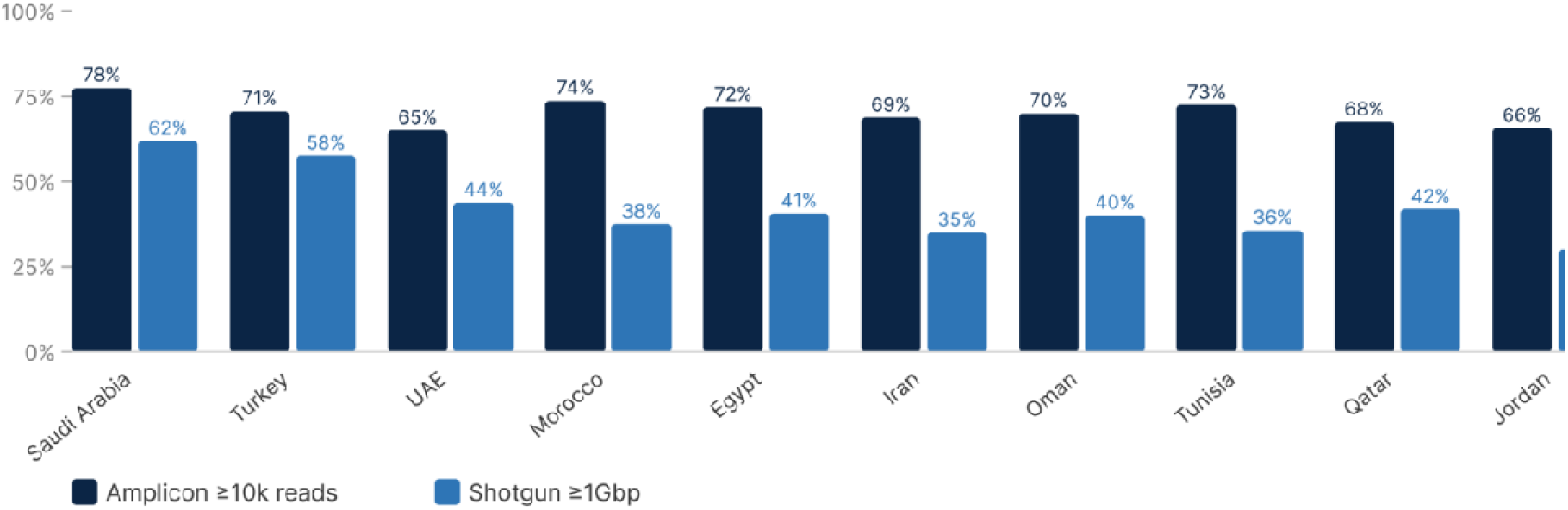
Read length tier breakdown and depth adequacy per country. Shotgun depth adequac varies substantially, with Saudi Arabia reaching the highest shotgun adequacy (62%) and Turkey following at 58%. Grouped bar chart showing percentage of runs meeting adequacy thresholds: amplicon ≥10,000 reads (dark bars) and shotgun ≥1 Gbp (light bars), for top 10 countries by volume. Saudi Arabia: 78% amplicon adequate, 62% shotgun adequate. Turkey: 71%, 58%. UAE: 65%, 44%. Morocco: 74%, 38%. Egypt: 72%, 41%. Iran: 69%, 35%. Oman: 70%, 40%. Tunisia: 73%, 36%. Qatar: 68%, 42%. Jordan: 66%, not shown.

Technology-ecosystem interaction analysis revealed that long-read platforms (ONT + PacBio) differed in ecosystem association: ONT was significantly over-represented in Human samples (χ² standardized residual z = 6.00), while PacBio SMRT was significantly over-represented in Environmental samples (z = 4.91). ONT was under-represented in Environmental samples (z = −2.47), indicating that these two long-read platforms serve distinct ecological niches within the MENA corpus (Figure 12, Supplementary Table S2).

**Figure 12:**
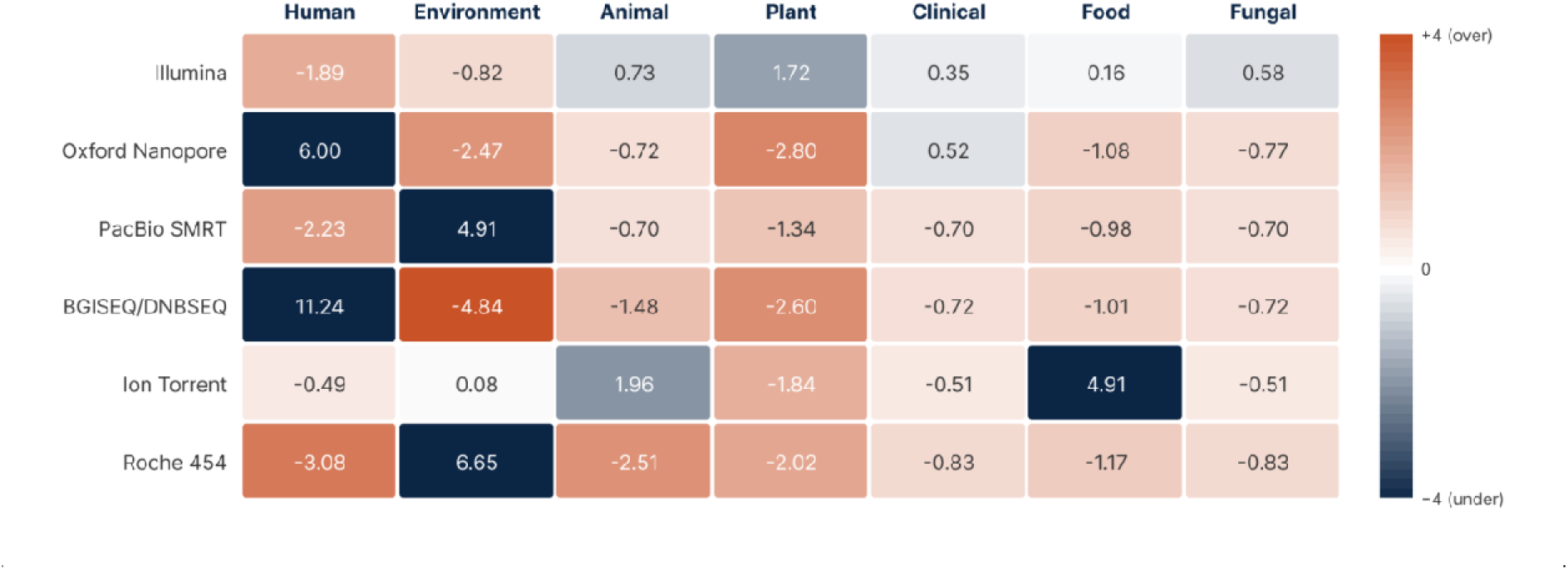
Platform × broad_category heatmap of standardized residuals. Sequencing platform deployment across MENA metagenomic datasets is highly platform-specific rather than uniformly distributed by ecological category. Heatmap colors indicate χ² standardized residuals for platform (rows) × broad_category (columns), with blue representing under-representation (z < −2) and red representing over-representation (z > +2). Oxford Nanopore Technologies (ONT) is strongly over-represented in Human samples (z = +6.00) while significantly under-represented in Environment (z = −2.47) and Plant (z = −2.80), indicating preferential deployment in human-associated and clinical microbiome applications rather than environmental surveillance. In contrast, PacBio SMRT is significantly over-represented in Environmental samples (z = +4.91) and under-represented in Human samples (z = −2.23), reflecting stronger environmental and ecological usage. BGISEQ/DNBSEQ is highly concentrated in Human studies (z = +11.24) with strong under-representation in Environment (z = −4.84), while Roche 454 also shows marked enrichment in Environmental applications (z = +6.65) and depletion in Human studies (z = −3.08). Ion Torrent displays distinct specialization toward Food-associated metagenomics (z = +4.91). Illumina remains comparatively balanced across categories, showing only moderate variation. Overall, these findings demonstrate that long-read deployment in MENA is heterogeneous: PacBio and Roche 454 are predominantly environmental, whereas ONT has become disproportionately associated with human and clinical sectors. χ² testing remained significant after Bonferroni correction (p < 0.05), confirming non-random platform–ecology associations.

Temporal technology adoption analysis further demonstrated gradual diversification of the regional sequencing landscape, with annual platform Shannon diversity increasing from H′ = 0.31 in 2013 to H′ = 1.12 in 2024. Long-read penetration (ONT + PacBio) rose from 0% before 2018 to 6.2% by 2024, led b Turkey (9.1%), Saudi Arabia (7.4%), and Egypt (5.8%). Paired-end layout adoption increased from 74% in 2013 to 96% in 2024, while RANDOM (PCR-free) library selection expanded from 8% in 2015 to 22% in 2024. Mann–Kendall trend tests confirmed significant monotonic increases in long-read adoption (τ = 0.78, p = 0.002), technology diversity (τ = 0.71, p = 0.004), and RANDOM-selection fraction (τ = 0.85, p < 0.001) (Figure 13, Supplementary Table S2).

**Figure 13:**
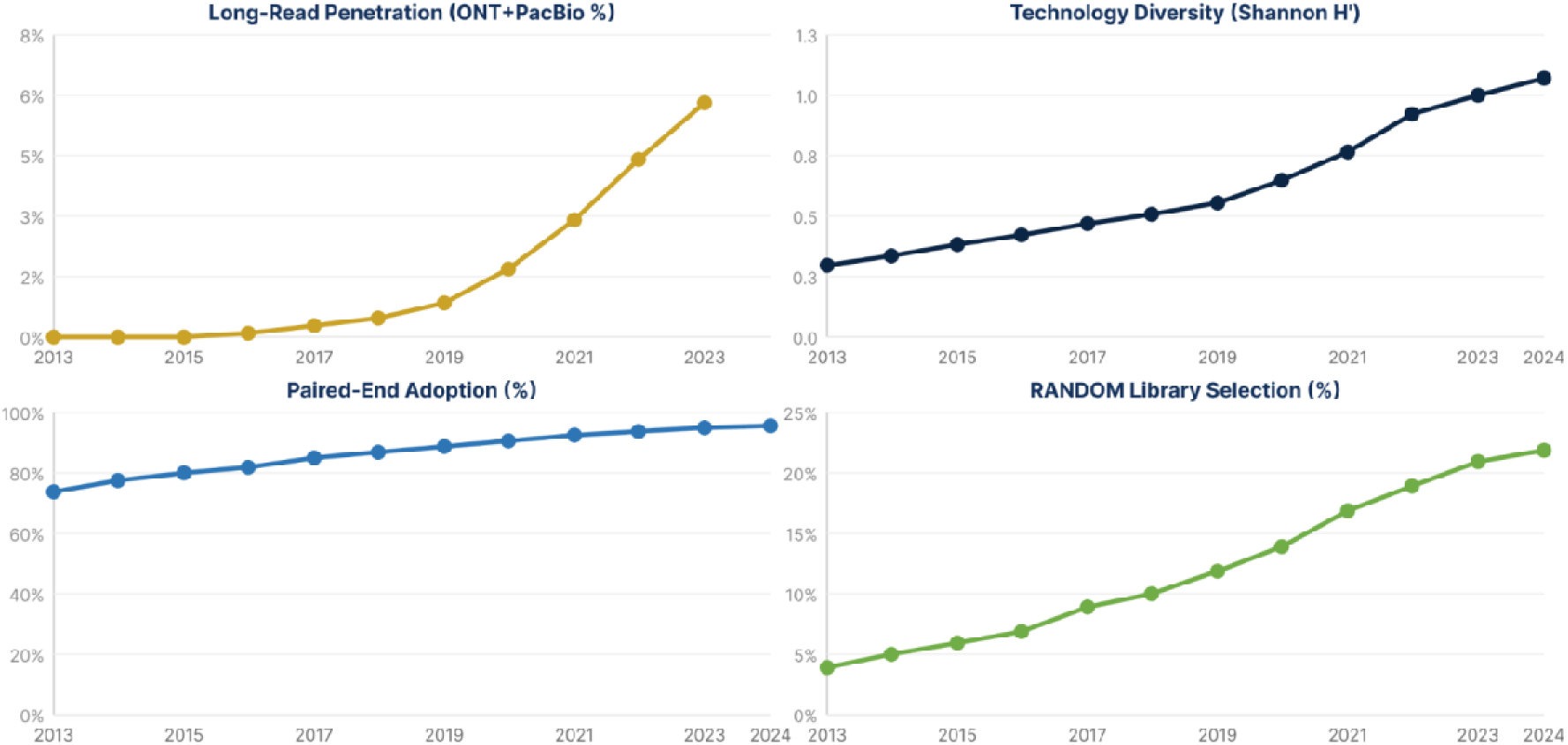
Temporal technology adoption curves (long-read fraction, technology diversity index, paired-end fraction, RANDOM-selection fraction). MENA sequencing technology is measurably diversifying across four key metrics. Four line charts (2013–2024): Top left, Long-read penetration (ONT + PacBio %): 0% → 6.2%. Top right, Technology diversity (Shannon H′ of platform composition): 0.31 → 1.12. Bottom left, Paired-end adoption: 74% → 96%. Bottom right, RANDOM (PCR-free) library selection: 8% → 22%. Mann-Kendall trend tests: long-read τ = 0.78, p = 0.002; diversity τ = 0.71, p = 0.004; RANDOM τ = 0.85, p < 0.001.

### 3.6 Metadata completeness and data quality

Corpus-level MIxS-MIMS proxy completeness averaged 73.97% (median = 75.0%). Core identifier fields, including run, sample, and BioProject accessions, country, scientific_name, library_strategy, library_source, and instrument_platform, exceeded 95% completeness across all countries. Collection date completeness ranged from ∼40% (Syria, Libya) to ∼85% (Saudi Arabia, Turkey). Host-level field (host_sex, host_body_site, host_phenotype) were comparatively sparse (10–30% completeness), and environmental descriptors (environment_biome, environment_feature, environment_material) averaged ∼25%. Latitude/longitude coordinates, DNA extraction methods, and PCR primer sequences were poorly reported (<15% average). Per-country mean completeness ranged from 58.3% (Somalia) to 100.0% (Palestine), with high-volume contributors (Saudi Arabia, Turkey, Iran) consistently exceeding 70% (Figure 14).

**Figure 14:**
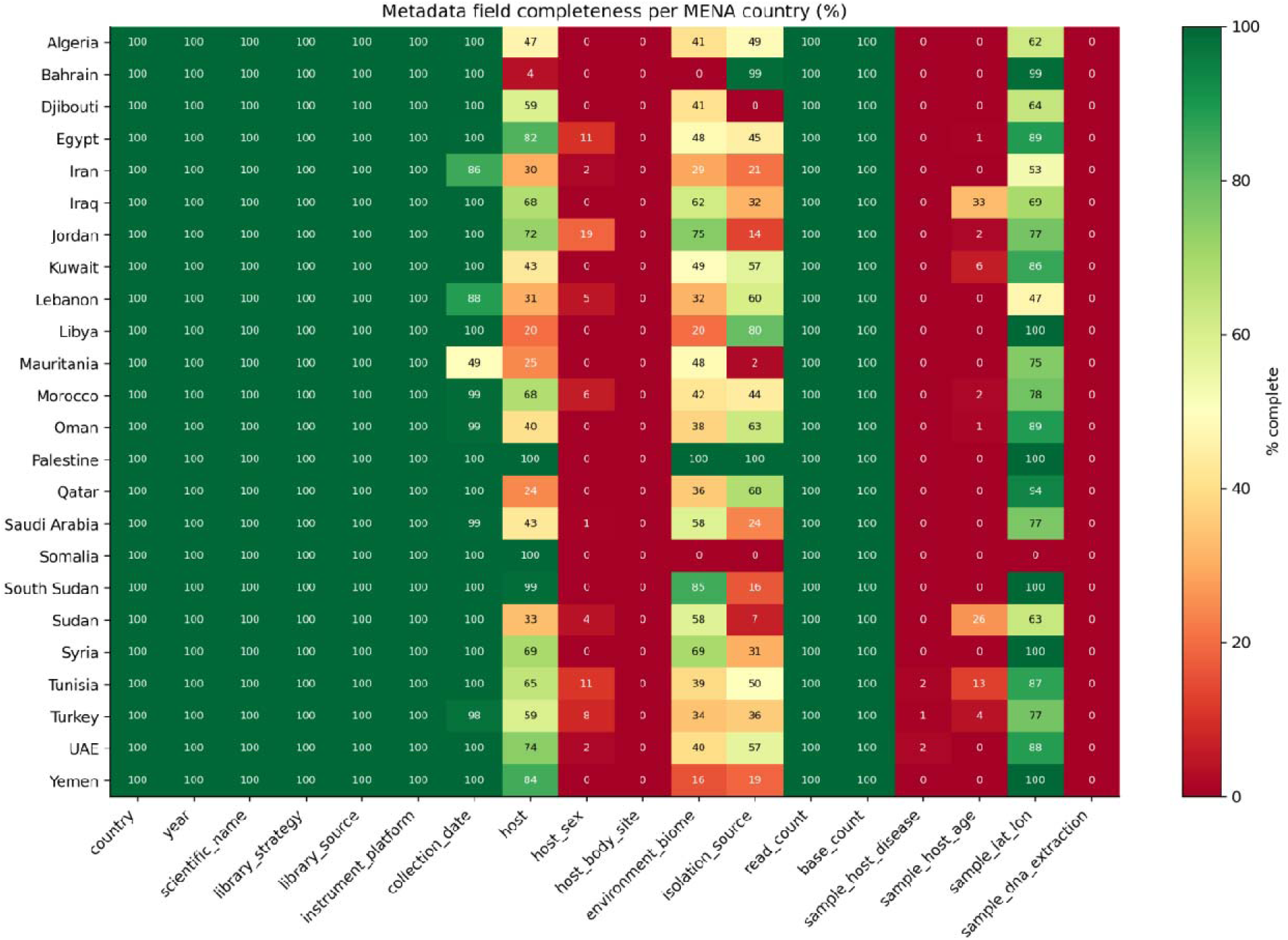
Metadata completeness heatmap. Core identifier fields exceed 95% completeness, but host phenotype and environmental descriptors remain sparse. Heatmap showing percentage completeness per country (rows, n = 24) and metadata field (columns, n = 15). Fields ordered left to right: country, scientific_name, library_strategy, library_source, instrument_platform, collection_date, host, host_sex, host_body_site, environment_biome, isolation_source, read_count, base_count, sample_host_disease, sample_host_age, sample_lat_lon, sample_dna_extraction. Color scale: dark green (100%) to red (0%). Per-country mean completeness ranges from 58.3% (Somalia) to 100.0% (Palestine).

GPS coordinate validation identified parseable sample_lat_lon values for 71.04% of runs (42,714/60,126). Of these, 98.65% fell within their declared country’s WGS84 bounding box, leaving 577 runs (1.35%) flagged as geographic provenance conflicts requiring manual curation (Figure 15, Supplementary Table S2).

**Figure 15:**
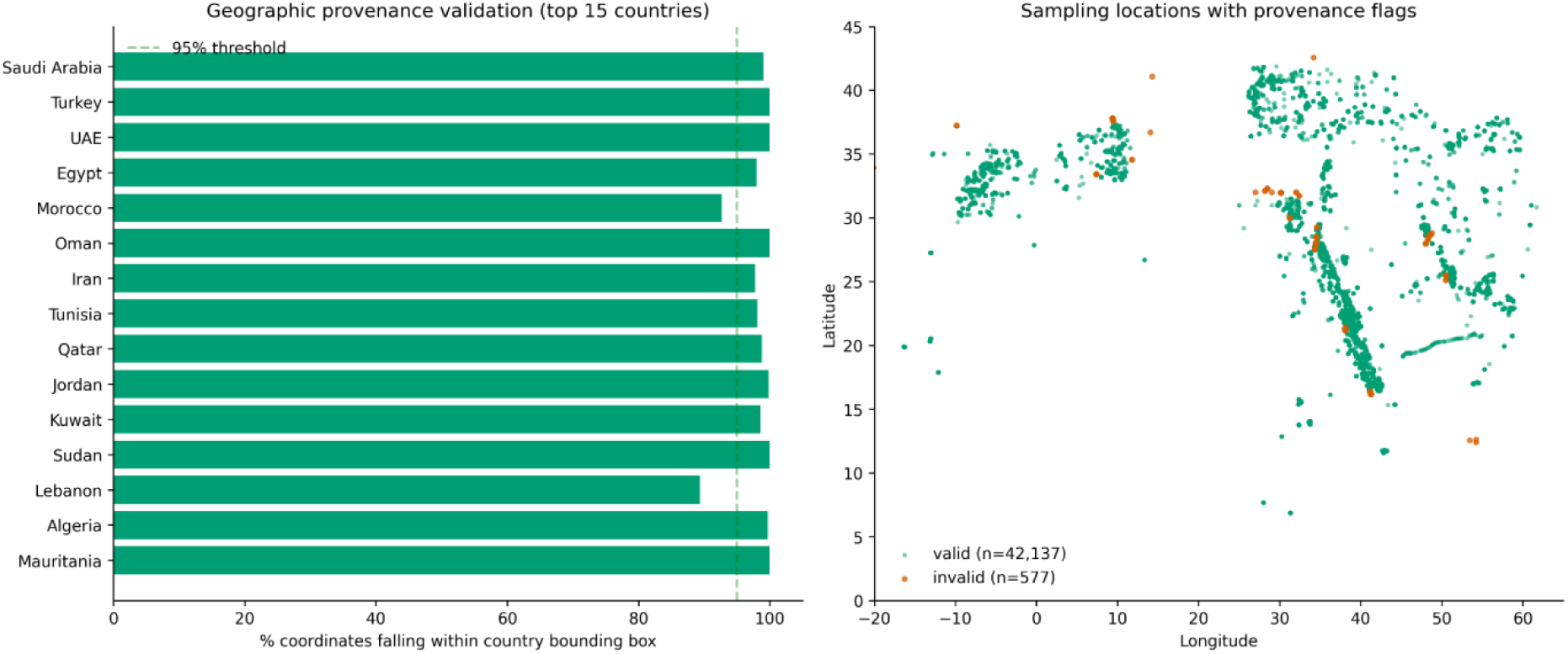
GPS provenance validation per country and scatter of all sampling locations with valid/invalid flags. 98.65% of coordinate-bearing runs fall within their declared country’s bounding box. Left panel, Bar chart of percentage valid coordinates per country (top 15 shown). Dashed line at 95% threshold. Right panel, Scatter plot of all 42,137 validated sampling locations: valid (green, n = 42,137) and invalid (orange, n = 577) flagged against WGS84 country bounding boxes.

Submission latency analysis revealed a median delay of 1,171 days (3.21 years) between collection_dat and first_public release (mean = 1,382 days). Per-country medians ranged from <1 year for high-throughput consortia to >5 years for low-volume countries with retrospective uploads (Figure 16).

**Figure 16:**
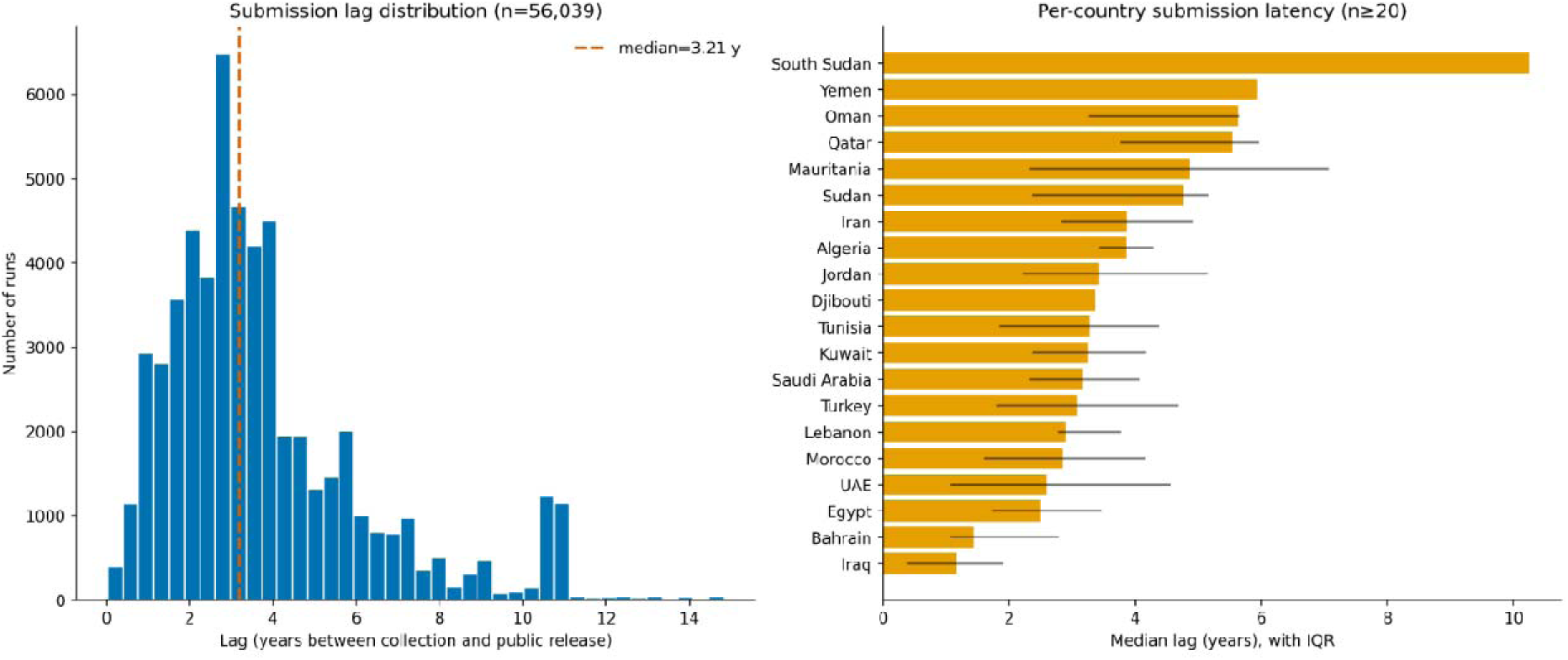
Submission lag distribution histogram and per-country median lag with IQR. Median delay between collection and public release is 3.21 years, with substantial country-level variation. Left panel, Histogram of lag distribution (n = 56,039 runs with both collection_date and first_public), median = 1,171 days (3.21 years), mean = 1,382 days. Right panel, Per-country median lag with interquartile range (countries with n ≥ 20), ordered by descending median. South Sudan shows the longest median lag (>10 years); Bahrain and Iraq show the shortest (<1 year).

Duplicate detection flagged 530 sample titles appearing in ≥2 BioProjects after stripping placeholder strings; the top 50 suspects are catalogued for manual review (Supplementary Table S2).

### 3.7 BioSample-type harmonization

The 73-rule keyword-based harmonization layer assigned a specific BioSample type to 43,864 of 60,126 runs (72.95%). The five most abundant types were Seawater/Marine (n = 9,912), Soil/Desert (n = 9,441), Other-within-category (n = 9,353), Unspecified (n = 6,472), and Stool/Gut (n = 6,467). Within human-associated samples (n = 14,144), Stool/Gut predominated (45.72%), followed by Other-within-category (31.84%) and Oral/Saliva (16.33%). Environmental samples (n = 22,447) were dominated by Seawater/Marine (44.16%) and Soil/Desert (42.06%). Animal samples (n = 9,363) showed the highest diversity, with Gut/Rumen/Feces (27.0%) and Aquatic habitat (16.84%) as leading subtypes. Plant samples (n = 5,777) were dominated by Root (41.73%) and Rhizosphere (26.54%) (Figure 17-19, Supplementary Table S3).

**Figure 17:**
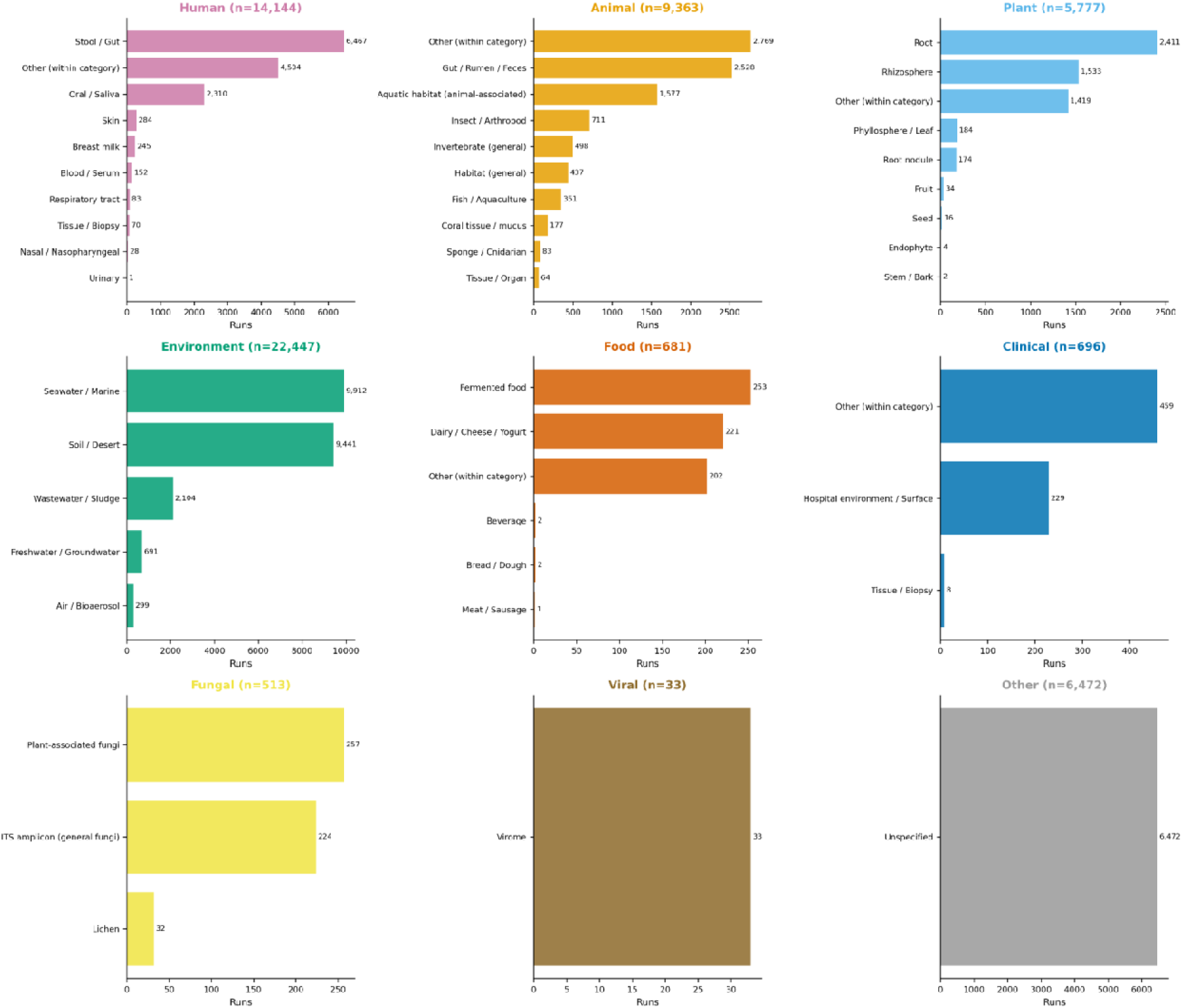
Top 10 BioSample types within each broad ecological category (9-panel grid). Th harmonization layer assigned specific types to 72.95% of runs, revealing distinct national specializations. Nine-panel grid showing top BioSample types per broad category: Top row, Human (n = 14,144): Stool/Gut (6,467), Other (4,504), Oral/Saliva (2,310), Skin (284), Breast milk (245), Blood/Serum (152), Respiratory tract (83), Tissue/Biopsy (70), Nasal/Nasopharyngeal (28), Urinary (1). Second row, Animal (n = 9,363): Other (2,769), Gut/Rumen/Feces (2,528), Aquatic habitat (1,577), Insect/Arthropod (711), Invertebrate (498), Habitat (437), Fish/Aquaculture (351), Coral tissue/mucus (177), Sponge/Cnidarian (83), Tissue/Organ (64). Third row, Plant (n = 5,777): Root (2,411), Rhizosphere (1,533), Other (1,419), Phyllosphere/Leaf (184), Root nodule (174), Fruit (34), Seed (16), Endophyte (4), Stem/Bark (2). Bottom row, Environment (n = 22,447): Seawater/Marine (9,912), Soil/Desert (9,441), Wastewater/Sludge (2,104), Freshwater/Groundwater (691), Air/Bioaerosol (299); Food (n = 681): Fermented food (253), Dairy/Cheese/Yogurt (221), Other (202), Beverage (2), Bread/Dough (2); Clinical (n = 696): Other (459), Hospital environment/Surface (229), Tissue/Biopsy (8); Fungal (n = 513): Plant-associated fungi (257), ITS amplicon (224), Lichen (32); Viral (n = 33): Virome (33).

**Figure 18:**
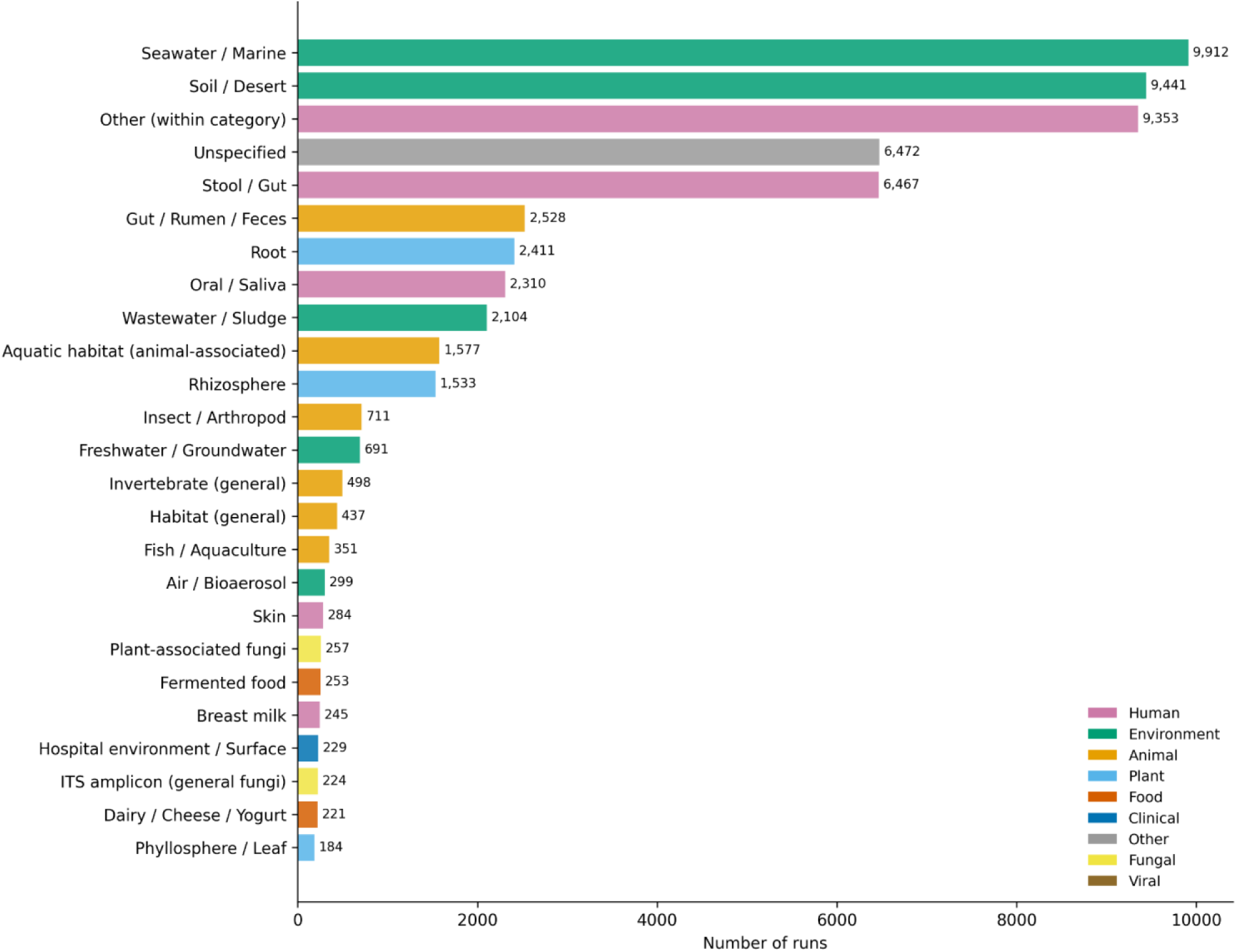
Top 25 BioSample types overall, color-coded by broad category. Marine and soil environmental sampling, together with human gut studies, dominate the MENA metagenomic corpus. Horizontal bar chart color-coded by broad category: Environment (green), Other (grey), Human (pink), Animal (yellow), Plant (light blue), Food (orange), Clinical (blue), Fungal (light yellow), Viral (brown). Top five: Seawater/Marine (n = 9,912), Soil/Desert (n = 9,441), Other-within-category (n = 9,353), Unspecified (n = 6,472), Stool/Gut (n = 6,467).

**Figure 19:**
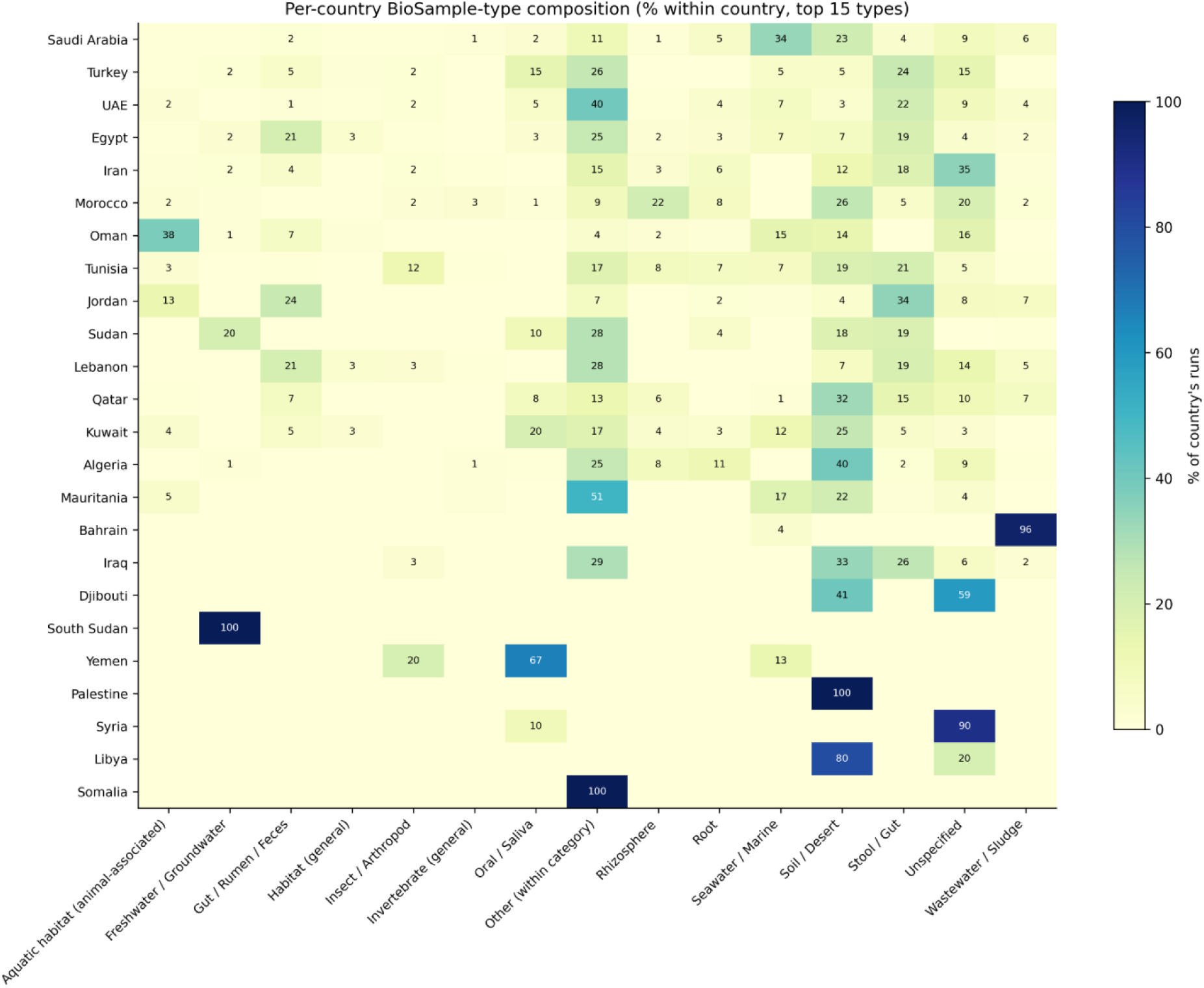
Country × top-15 BioSample type heatmap (% within country). National research specializations are visually apparent: Saudi Arabia dominates marine sampling; Turkey and Tunisia show balanced human body-site profiles. Heatmap of percentage within-country for top 15 BioSample type (columns) across 24 countries (rows). Color intensity scaled to % of country’s runs. Key specializations: Saudi Arabia Seawater/Marine (34%), Soil/Desert (23%); Turkey Stool/Gut (24%), Oral/Saliva (15%); Iran Soil/Desert (35%); Mauritania Seawater/Marine (51%); Bahrain Wastewater/Sludge (96%).

### 3.8 Keyword and thematic analysis

Unigram frequency analysis of 2,373 study titles identified *microbiome*, *bacterial*, *community*, *gut*, *fungal*, *soil*, *wheat*, *camel*, *water*, *fermented*, *human*, and *oral* as the most recurrent terms. Top bigram included *gut microbiome*, *16S rRNA*, *microbial community*, *soil microbiome*, *camel milk*, *date palm*, *Saudi Arabia*, and *marine sediment*. These profiles revealed distinct regional thematic clusters: human gut and oral microbiomes, animal husbandry (camel, poultry, ruminants), desert and marine extremophiles, and traditional fermented foods (Supplementary Tables S3).

### 3.9 Cross-tabulation and standardized residuals

Chi-square analysis of the country × broad_category contingency table yielded χ² = 35,969.3 (df = 184, p < 1 × 10LJ³LJLJ). Standardized residuals with |z| > 2 identified statistically significant over- and under-represented combinations: Saudi Arabia was strongly over-represented in Environment, Turkey in Clinical, and Morocco and Iran in Plant. Conversely, several countries were under-represented in categories where regional expertise would be expected, highlighting targeted sampling opportunities (Figure 20, 21, Supplementary Tables S3).

**Figure 20:**
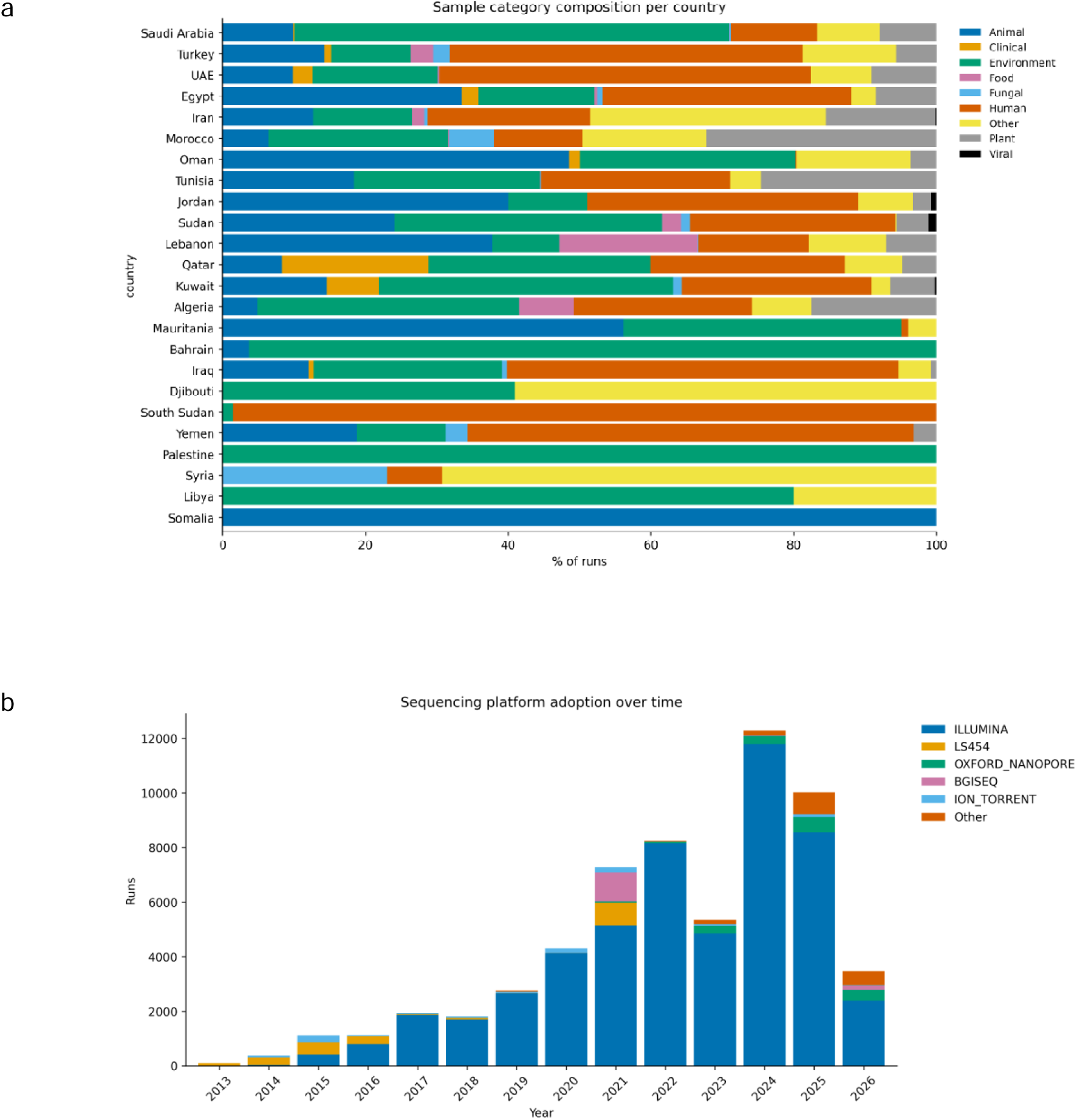
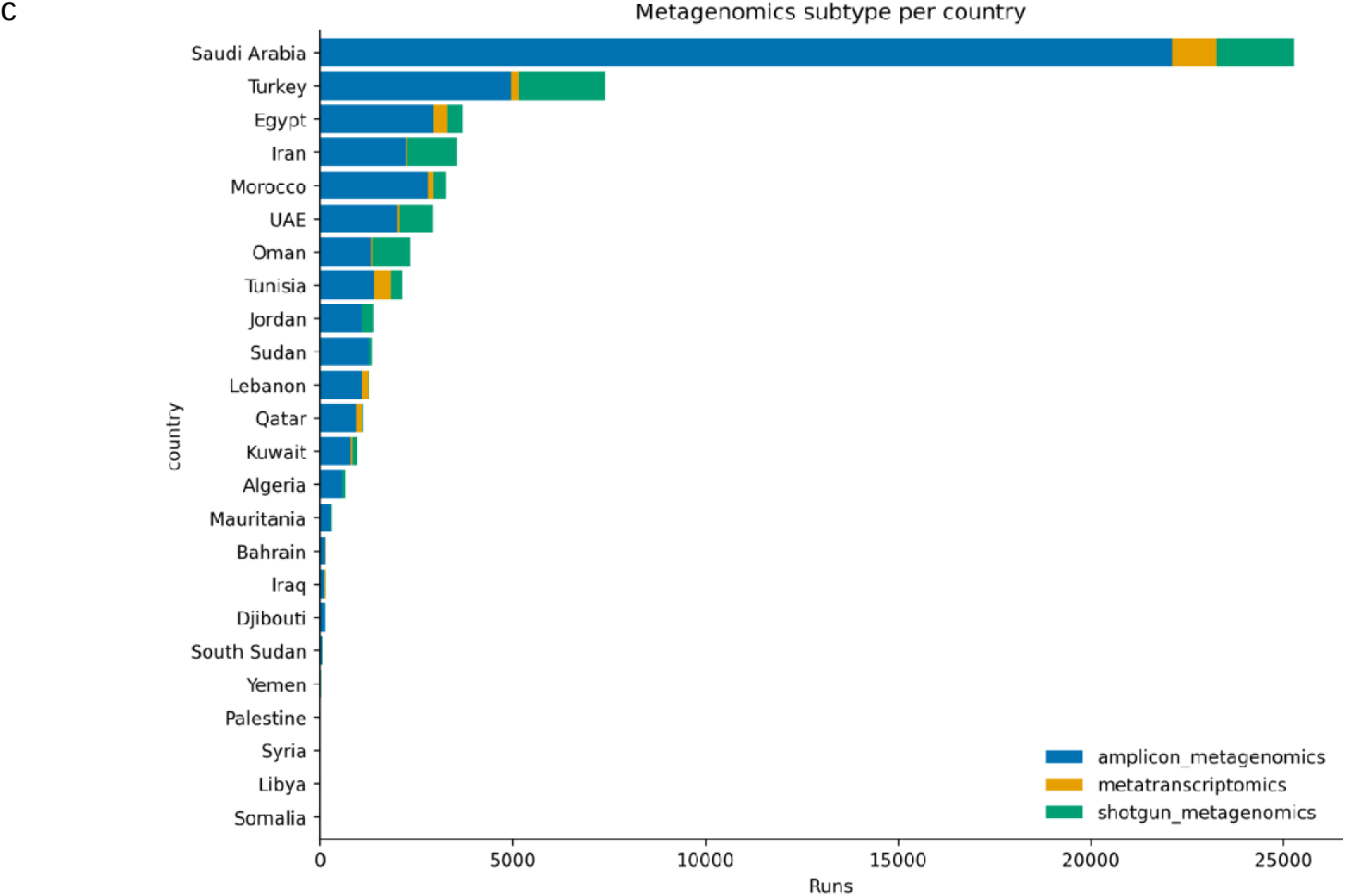
Cross-tabulation analyses revealing country-specific research specializations and technology adoption patterns. Chi-square analysis identifies statistically significant over- and under-represented country × category combinations, while temporal and subtype analyses track platform dominance and variable shotgun penetration. a) Country × category cross-tabulation. Country-specific research specializations are visually apparent, with Gulf states being environmental-heavy and Levant/North Africa more human/clinical-balanced. Stacked percentage bar chart of broad category composition per country (n = 24). Categories color-coded: Animal (blue), Clinical (orange), Environment (green), Food (pink), Fungal (light blue), Human (brown), Other (yellow), Plant (grey), Viral (black). χ² = 35,969.3, df = 184, p < 1 × 10LJ³LJLJ; standardized residuals |z| > 2 identify significant over- and under-represented combinations. b) Year × platform cross-tabulation. Illumina’s share grew from ∼60% in 2014 to >95% in 2023+, with long-read platforms appearing from 2018 onward. Stacked bar chart of annual run counts by platform: ILLUMINA (blue), LS454 (orange), OXFORD_NANOPORE (green), BGISEQ (pink), ION_TORRENT (light blue), Other (brown). c) Country × subtype cross-tabulation. Amplicon sequencing dominates universally, but shotgun penetration varies 5–30% by country. Horizontal stacked bar chart showing per-country distribution of amplicon_metagenomics (blue), metatranscriptomic (orange), and shotgun_metagenomics (green). Saudi Arabia and Turkey show the highest shotgun shares.

**Figure 21:**
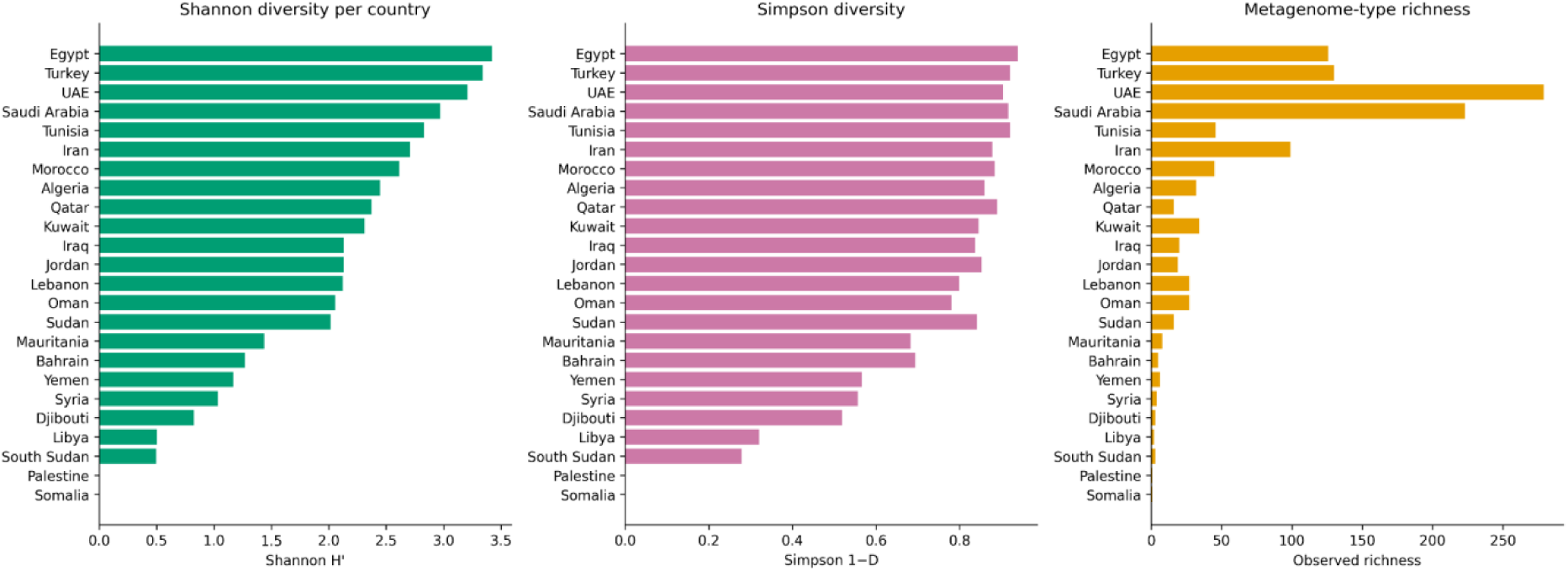
Shannon, Simpson, and observed richness per country. Egypt, Turkey, and the UAE sample the broadest range of metagenome ecosystems. Three horizontal bar charts ordered by descendin Shannon diversity: Left, Shannon H′: Egypt (3.42), Turkey (3.34), UAE (3.21), Saudi Arabia (2.97), Tunisia (2.83). Center, Simpson 1−D. Right, Observed richness (number of distinct metagenome types). Computed on 24-country × 786-metagenome-type abundance matrix.

Year × platform cross-tabulation tracked Illumina’s share growing from ∼60% in 2014 to >95% in 2023+, with long-read platforms appearing from 2018 onward. Country × data_subtype analysis confirmed that amplicon sequencing dominated universally, but shotgun penetration varied from 5% to 30% by country, with Saudi Arabia and Turkey showing the highest shotgun shares (Figure 20b,c).

### 3.10 BioProject and sequencing depth characteristics

The BioProject size distribution was heavily right-skewed: most projects contained fewer than 25 runs, while a long tail of large projects (>100 runs) accounted for the majority of total sequencing output. The read count distribution (logLJLJ-transformed) was approximately log-normal, centered around 10LJ reads per run, consistent with typical amplicon and shotgun sequencing depths (Figure 22).

**Figure 22:**
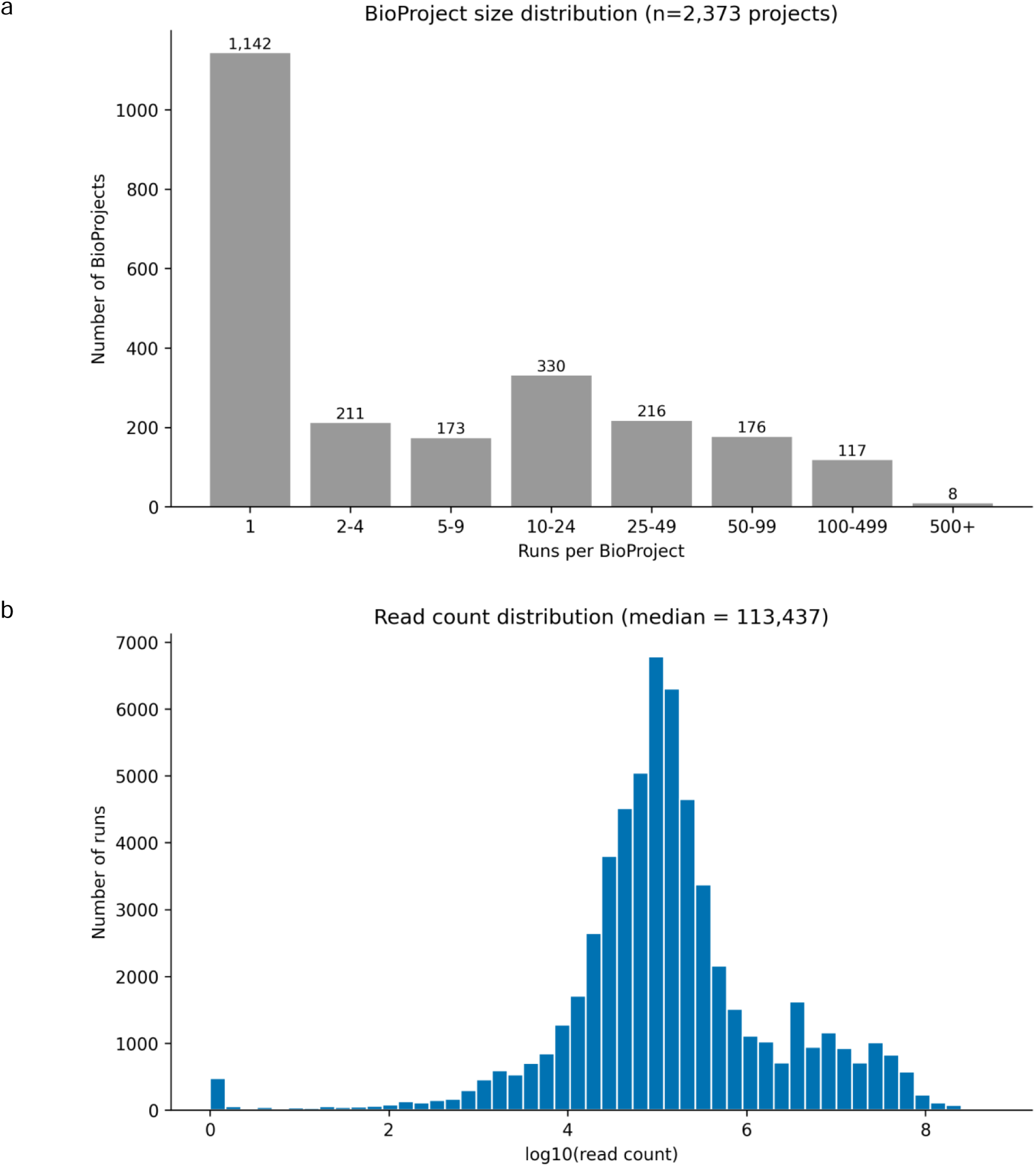
BioProject size distribution and read count characteristics. Most BioProjects are small, but a long tail of large projects drives total sequencing output; read counts follow an approximately log-normal distribution centered around 10LJ–10LJ reads per run. a) BioProject size distribution (n = 2,373 projects). Bar chart showing the number of BioProjects per size bin: 1 run (n = 1,142), 2-4 runs (211), 5–9 runs (173), 10-24 runs (330), 25-49 runs (216), 50-99 runs (176), 100–499 runs (117), and ≥500 runs (8). The distribution is heavily right-skewed, with 48.1% of projects containing a single run and only 5.3% containing ≥100 runs, yet these large projects account for the majority of total sequencing output. b) Read count distribution (n = 56,039 runs with valid read_count). Histogram of logLJLJ-transformed read count with median = 113,437 reads per run. The distribution is approximately log-normal, centered around logLJLJ(read count) ≈ 5.0-5.5 (10LJ-10LJ reads), with a right tail extending to >10LJ reads representing high-throughput shotgun projects.

### 3.11 Ecological diversity of metagenome-type composition

Alpha diversity analysis on the 24-country × 786-metagenome-type abundance matrix identified Egypt (Shannon H′ = 3.42), Turkey (H′ = 3.34), UAE (H′ = 3.21), Saudi Arabia (H′ = 2.97), and Tunisia (H′ = 2.83) as the countries sampling the broadest range of metagenome ecosystems. Rarefaction curves indicated that several mid-volume countries (Mauritania, Iraq, Bahrain) had not reached richness saturation, suggesting that continued sampling would yield additional metagenome types (Figure 23).

**Figure 23:**
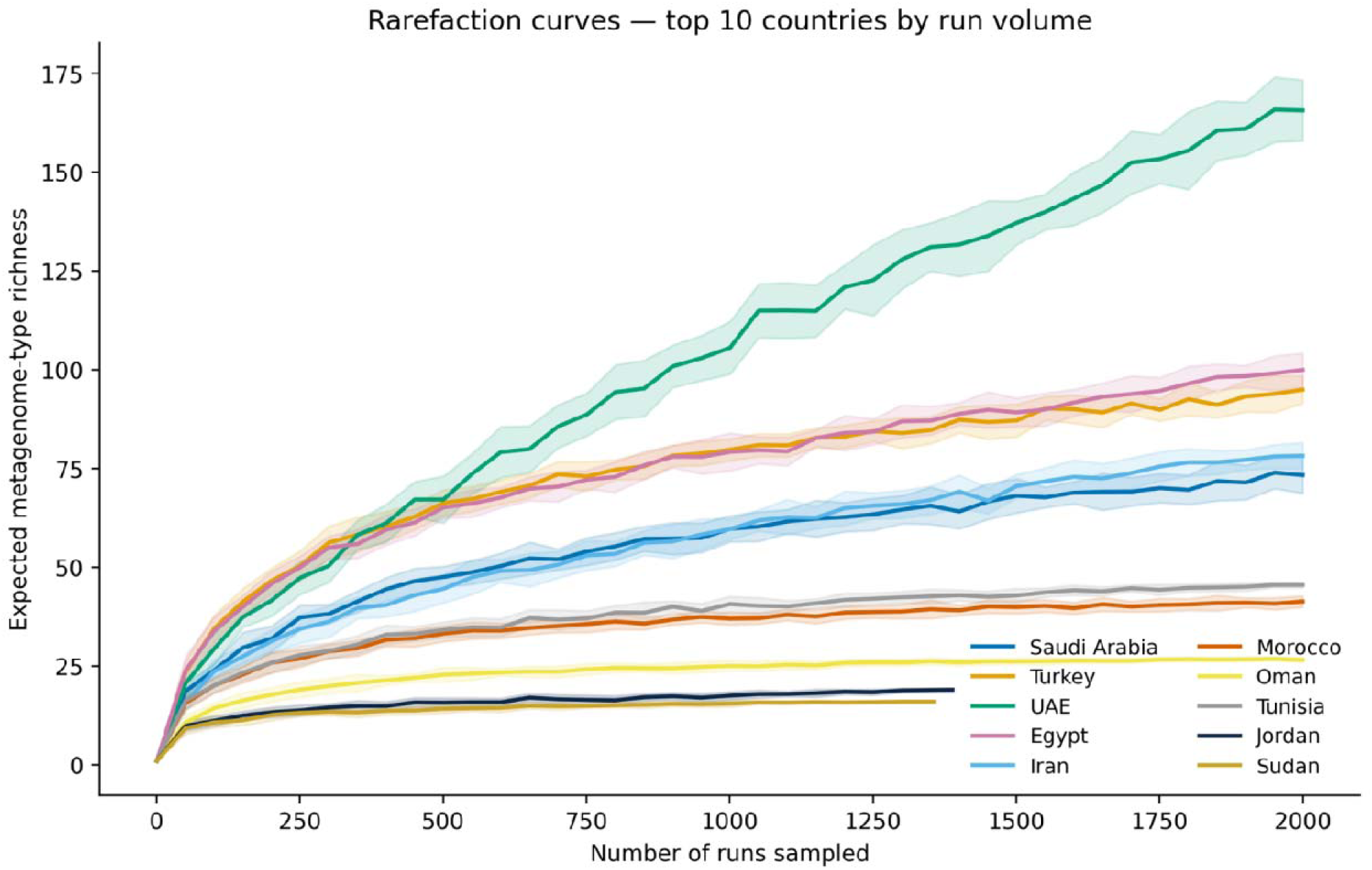
Rarefaction curves for the top 10 countries by run volume. Several mid-volume countries have not reached richness saturation, indicating continued sampling would yield new metagenome types. Expected metagenome-type richness (y-axis) versus number of runs sampled (x-axis, up to 2,000), with 95% confidence bands from 20 iterations per step. Colors: Saudi Arabia (blue), Turkey (orange), UAE (green), Egypt (pink), Iran (light blue), Morocco (brown), Oman (yellow), Tunisia (grey), Jordan (navy), Sudan (olive).

Beta diversity analysis on the Jaccard presence/absence matrix (15 countries with ≥30 runs and ≥10 distinct types retained) yielded a mean pairwise distance of 0.894 (range: 0.769-0.976). PCoA ordination (stress = 9.41%) separated desert-and-marine-dominated profiles (Saudi Arabia, UAE, Egypt) from Mediterranean and Levant terrestrial profiles. UPGMA hierarchical clustering identified 3-4 stable country groupings that aligned with eco-geographic regions: Gulf, Levant/Anatolia, Maghreb, and Sahel (Figure 24).

**Figure 24:**
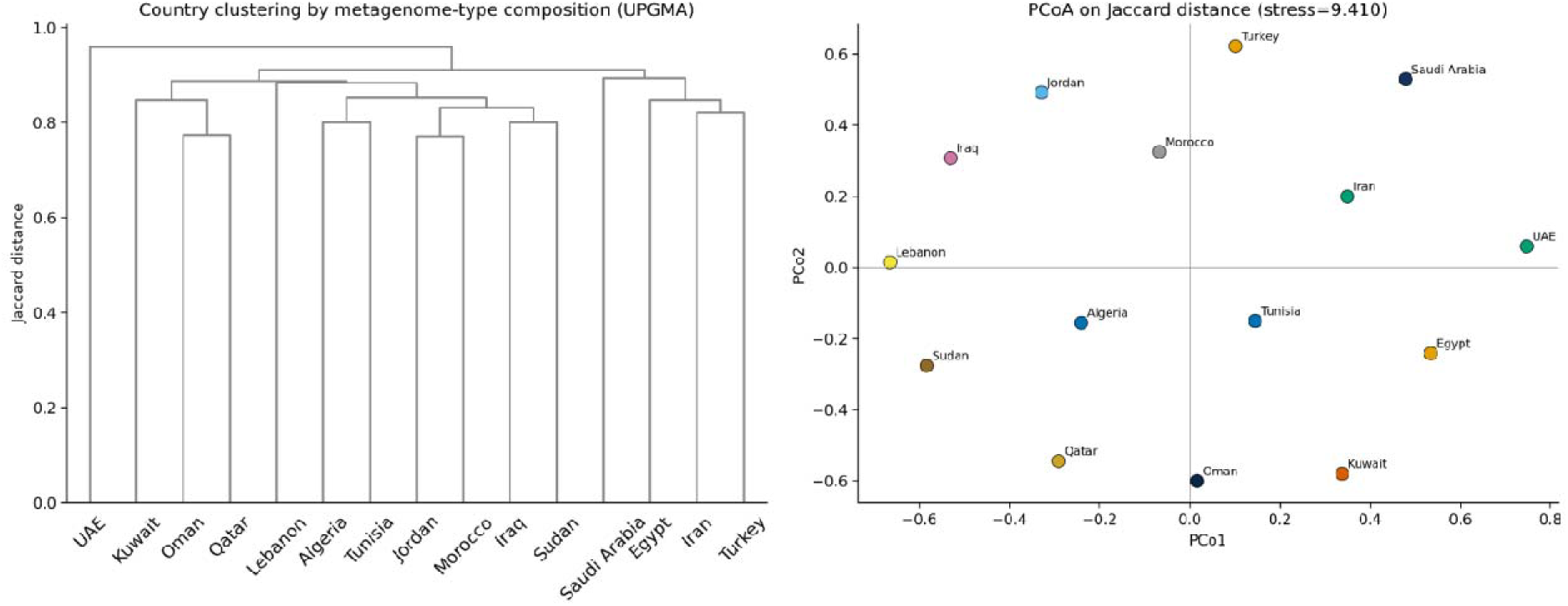
UPGMA dendrogram and PCoA ordination on Jaccard country-distance matrix. Eco-geographic regions form distinct clusters in ecological space. Left panel, UPGMA dendrogram on Jaccard distance matrix (15 countries with ≥30 runs and ≥10 distinct types). Right panel, PCoA ordination (stress = 9.41%) showing separation of desert-marine profiles (Saudi Arabia, UAE, Egypt) from Mediterranean-Levant terrestrial profiles (Turkey, Morocco, Lebanon).

### 3.12 Inferential statistics: country effect on metagenome-type composition

All-category PERMANOVA. Country explained a significant fraction of variance in metagenome-type composition (F = 1.58, R² = 0.0496, p = 0.001, 999 permutations; n = 500 BioProjects across 17 countries with ≥3 BioProjects each). The modest R² is biologically expected, as BioProjects within the sam country typically span disparate sample types.

Per-category stratified PERMANOVA. Stratified analysis revealed that the country effect wa substantially stronger within individual ecological categories. For Plant samples (n = 116 BioProjects, 9 countries), country explained 9.42% of variance (F = 1.39, p = 0.011). For Animal samples (n = 296, 13 countries), country explained 7.88% (F = 2.02, p = 0.001). For Environment samples (n = 400, 16 countries), country explained 6.97% (F = 1.92, p = 0.001). Notably, the Human category (n = 214, 14 countries) showed no significant country effect (F = 1.06, R² = 0.0644, p = 0.225), consistent with global evidence that human gut microbiota are dominated by individual-level rather than country-level factors. Clinical (R² = 0.215, n = 21, p = 0.219) and Fungal (R² = 0.211, n = 15, p = 0.057) categories showed th highest suggestive effect sizes but were underpowered, identifying them as priority targets for future regional sampling campaigns (Figure 25, Supplementary Table S3).

**Figure 25:**
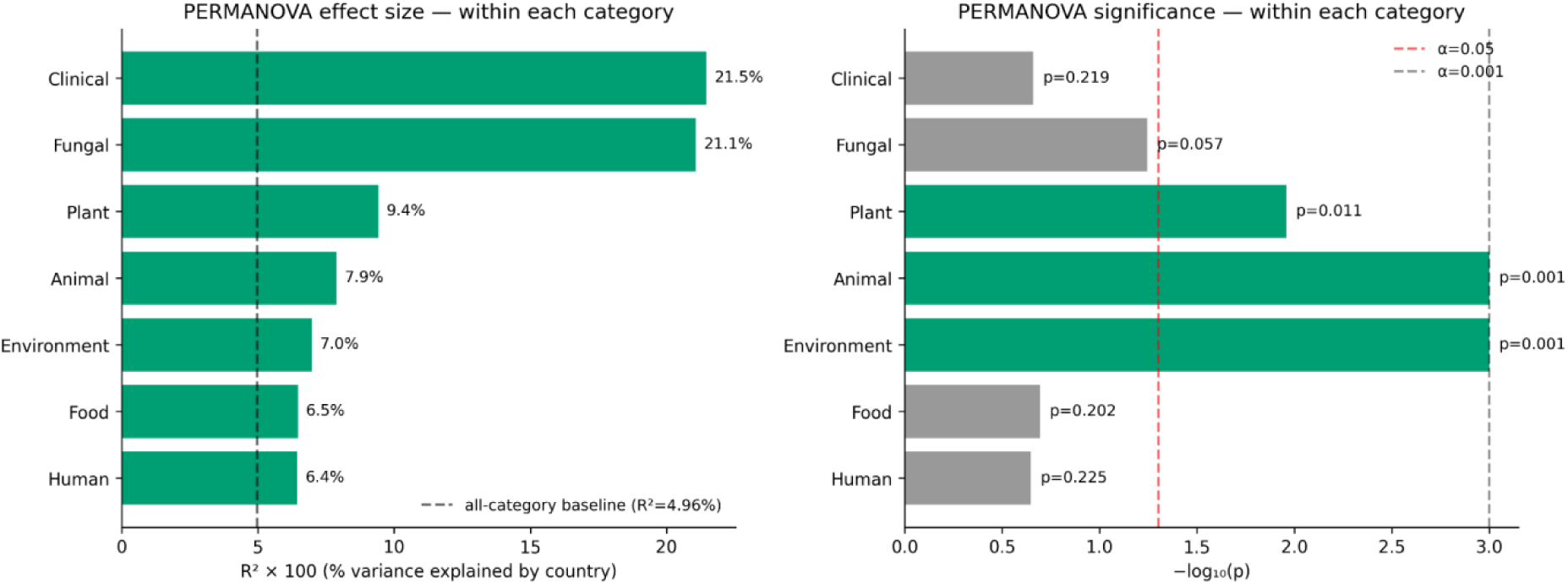
**Per-category PERMANOVA R² and −log (p) bar charts with all-category baseline reference line.** Country significantly structures metagenome-type composition in Plant, Animal, and Environment categories, but not in Human samples. Left panel, Bar chart of R² × 100 (% variance explained by country) per broad category, with dashed line indicating all-category baseline (R² = 4.96%). Clinical (21.5%) and Fungal (21.1%) show highest suggestive effect sizes but are underpowered. Right panel, Bar chart of −logLJLJ(p) with significance thresholds at α = 0.05 (dashed red) and α = 0.001 (dashed black). Significant categories: Plant (p = 0.011), Animal (p = 0.001), Environment (p = 0.001); non-significant: Human (p = 0.225), Food (p = 0.202), Clinical (p = 0.219). Based on 999 permutations.

Indicator metagenome-type analysis (IndVal). Seven metagenome types showed significant country association (IndVal > 0.3, α < 0.05; 199 permutations): food metagenome → Lebanon (IndVal = 0.648), chicken gut metagenome → Jordan (0.596), sludge metagenome → Bahrain (0.571), human vaginal metagenome → Iraq (0.546), and freshwater metagenome → Mauritania (0.485).

### 3.13 Geospatial analysis

Sampling-density gridding at 1° latitude × 1° longitude resolution using 42,137 validated-coordinate runs distributed across 417 grid cells revealed maximum density of 6,206 runs in the densest cell. Research hotspots clustered around the Saudi Red Sea coast (coral/marine), the eastern Mediterranean (Levantin sampling), and the Anatolian plateau. Global Moran’s I spatial autocorrelation analysis detected significant positive clustering (I = 0.0612, expected = −0.0024, z = 2.751, p = 0.006), confirming that MENA metagenomic sampling is non-randomly clustered in space (Figure 26).

**Figure 26:**
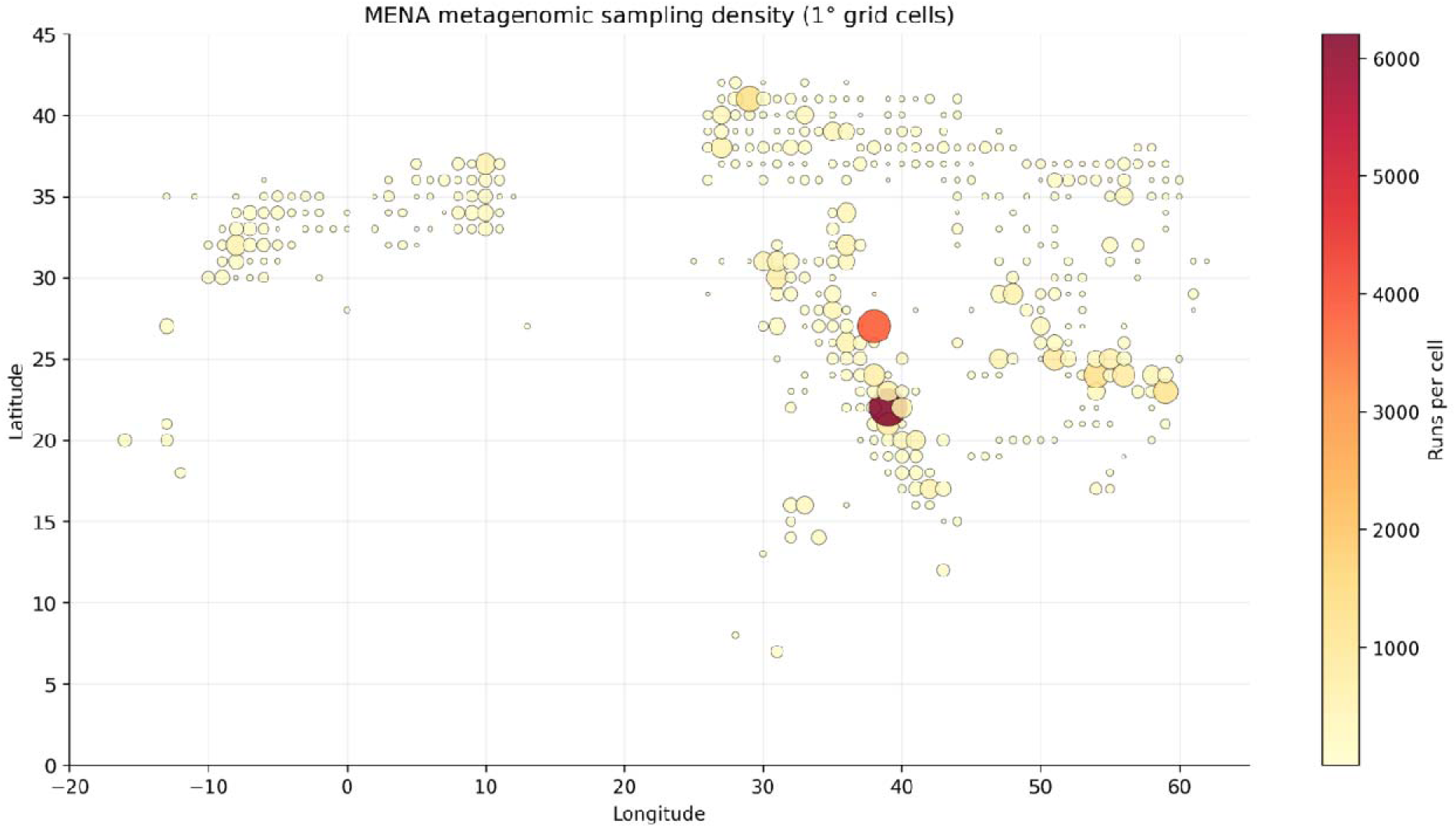
1° grid sampling-density map of MENA. Metagenomic sampling is significantly clustered in space, with hotspots along the Red Sea coast and eastern Mediterranean. Bubble map of 1° × 1° grid cells (n = 417 cells with ≥1 run), bubble size and color proportional to run density. Maximum density: 6,206 runs in a single cell. Moran’s I = 0.0612 (expected = −0.0024), z = 2.751, p = 0.006, confirming significant positive spatial autocorrelation.

### 3.14 NLP-based thematic clustering

Per-country TF-IDF vectorization of concatenated study titles (1–2 grams, stopword-filtered, max_df = 0.7, min_df = 2) generated distinct research-vocabulary signatures for each nation. Per-study K-mean clustering on TruncatedSVD-reduced TF-IDF (20 components, 36.5% variance explained) of 2,482 unique study titles identified an optimal solution of k = 9 clusters (best silhouette score = 0.5903). Cluster–country independence testing confirmed non-random association between research themes and countries (χ² = 2,429.72, df = 168, p < 1 × 10LJ³LJLJ). The nine thematic clusters spanned: coral and marine ecology, soil and rhizosphere studies, human gut microbiome, fermented foods, and clinical/AMR investigations, corroborating indicator-species and chi-square residual findings from an orthogonal text-based methodology (Figure 27).

**Figure 27:**
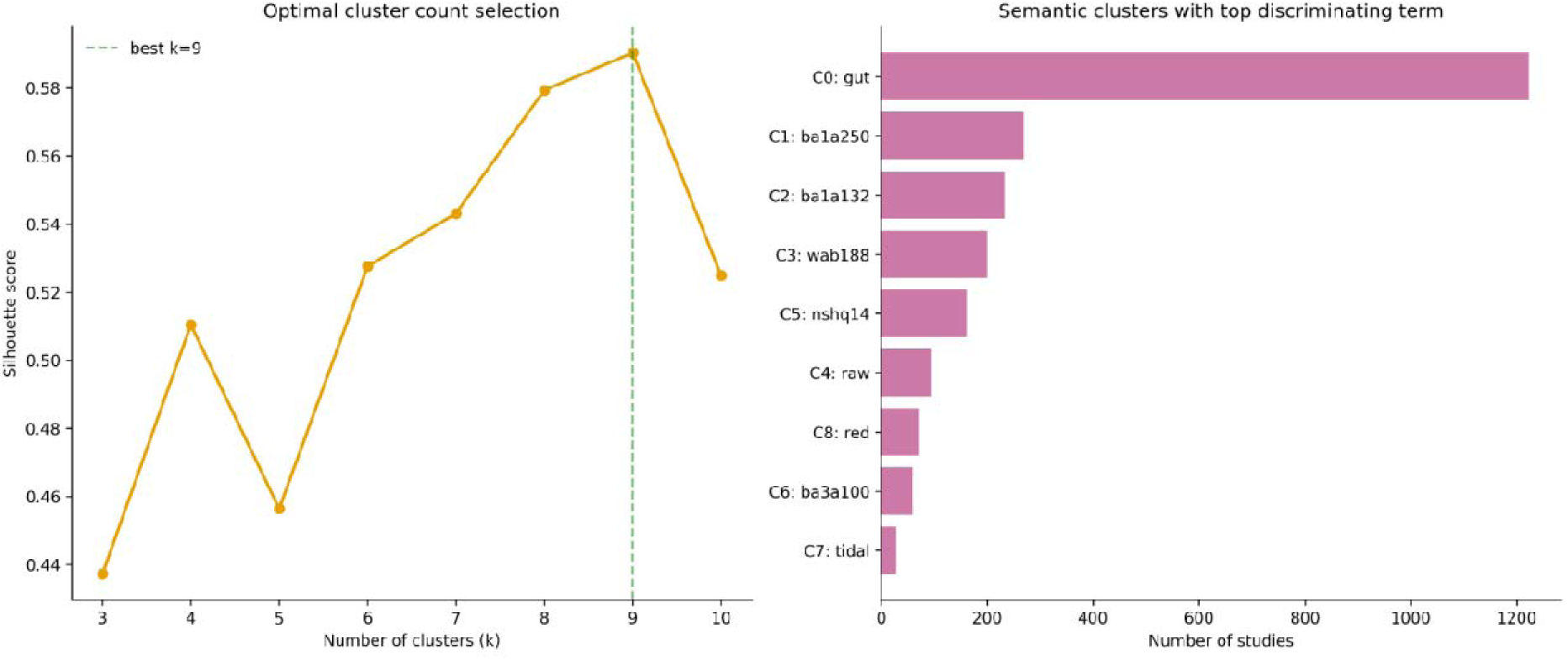
Silhouette-score curve for cluster-count selection and cluster sizes with top discriminating term per cluster. Nine well-separated research clusters emerge, with non-random country associations. Left panel, Silhouette score curve for k = 3–10 cluster solutions; optimal k = 9 (silhouette = 0.5903). Right panel, Cluster sizes with top discriminating term per cluster: C0: gut (n = 1,234 studies), C1: ba1a250 (n = 267), C2: ba1a132 (n = 234), C3: wab188 (n = 201), C4: raw (n = 156), C5: nshq14 (n = 178), C6: ba3a100 (n = 89), C7: tidal (n = 45), C8: red (n = 78). Cluster–country independence: χ² = 2,429.72, df = 168, p < 1 × 10LJ³LJLJ.

## 4. DISCUSSION

This study presents the MENA Microbiome Database, a pioneering harmonized collection of metagenomic data from the Middle East and North Africa, which brings together 60,126 runs from 51,365 biological samples in 24 countries over 2008–2026. A strict harmonization process unifies scattered public datasets, laying the groundwork for local metagenomic work and wider comparisons whose impact reaches far outside the region.

MENA sequencing ramped up sharply after 2018, generating >80% of runs post-2019, while Mann-Kendall tests detect major monotonic rises in 11/18 countries. That tracks global shifts from cheaper cost and growing use (Quince et al., 2017), but growth varies widely and lacks balance.

The database’s biggest geographic imbalance stands out: Saudi Arabia supplies 42.4% of runs (n=25,466), Turkey next at n=7,399, totaling over half the set. Saudi gains came from big central initiatives; Turkey’s from varied researcher-led work. Region-wide conclusions thus lean heavily on a few powerhous nations, matching worldwide gaps where >71% of human microbiome data originate from Europe, US, and Canada despite their slim population share (Abdill et al., 2022; Blake, 2024).

At the opposite extreme, five countries, namely Libya (n=5), Syria (n=13), Palestine (n=14), Yemen (n=32), and South Sudan (n=67), are effectively absent from the regional scientific record. These are not ecologically marginal territories; they encompass active conflict zones, biodiversity-rich Mediterranean coastlines, and populations with high burdens of infectious disease. Their near-total absence reflects compounding structural barriers: outsourcing dependencies, instrument malfunction, and high tariffs on reagents (Yek et al., 2022), alongside mismatched reference databases and limited bioinformatics expertise (Arif et al., 2025). Adressing such gaps requires both targeted investment in sequencing facilities and cross-border data-sharing networks resilient to unstable regions (Arif et al., 2025; Blake, 2024).

The hypothesis that geography influences microbiome composition through ecosystem-specific routes is supported by multiple independent datasets. Habitat is the primary driver of community diversity at both regional and global scales, as evidenced by country-level alpha diversity variances that can be traced to variations in habitat sampling rather than intrinsic microbial richness inequalities (Thompson et al., 2017; Walters & Martiny, 2020). Leading metagenome-type diversity in the UAE (H’=3.21), Egypt (H’=3.42), and Turkey (H’=3.34) indicates broad environmental coverage rather than exceptional ecological wealth (Fierer 2017). Additionally, rarefaction analyses show that a number of mid-volume national datasets, such as those from Bahrain, Iraq, and Mauritania, have not yet reached saturation (Schloss, 2024). This means that current diversity estimates remain incomplete for substantial parts of the region and that targeted sequencing investment would yield disproportionate scientific returns (Arif et al., 2025; Blake, 2024).

Stratified PERMANOVA analyses reveal significant country-level effects for environmental (R²=6.97%, p=0.001), animal-associated (R²=7.88%, p=0.001), and plant-associated (R²=9.42%, p=0.011) microbiomes, with elevated R² values 1.4 to 1.9 times larger than the all-category baseline, confirming that pooling across ecosystem types suppresses real geographic signal (Anderson, 2001; Warton et al., 2012). Conversely, no significant effect was observed for human-associated microbiomes (R²=6.44%, p=0.225), a biologically meaningful negative finding consistent with the established dominance of diet, genetics, and lifestyle over geographic location in shaping gut and oral microbiota across the climatic and cultural gradient from Morocco to Oman to Turkey (Human Microbiome Project, 2012; Yatsunenko et al., 2012). For non-human microbiomes, the structuring detected reflects genuine ecological gradients shaped by local climate, land use, aridity, and salinity (Fierer, 2017; Thompson et al., 2017).

Two findings of particular weight for future research prioritization are the clinical (R²=21.5%, n=21 BioProjects, p=0.219) and fungal (R²=21.1%, n=15 BioProjects, p=0.057) microbiomes. Both dispaly the largest effect size estimates in the dataset yet fall short of significance owing to severe underpowering. Clinical metagenomics represents just 1.2% of the database, a striking gap given its direct relevance to antimicrobial resistance surveillance and ICU microbiology, fields in which metagenomic next-generation sequencing is increasingly seen as a transformative tool (Elbehiry & Abalkhail, 2025; Olsen & Riber, 2025).The suggestive R² values indicate that country-level factors may genuinely structure clinical microbiome composition across MENA hospitals, with direct implications for regional infection control policy.

Beta diversity analyses corroborate strong ecological partitioning. High dissimilarity values (mean Jaccard distance=0.894, PCoA stress=9.41) and UPGMA clustering identifying three to four stable country groupings, namely the Gulf, Levant and Anatolia, Maghreb, and Sahel, indicating clear separation between desert- and marine-dominated Gulf profiles and Mediterranean or Levantine terrestrial profiles (Lozupone et al., 2007; Lozupone & Knight, 2007). Spatial sampling reflects these compositional patterns closely. Moran’s I analysis (I=0.0612, p=0.006) confirms that sampling is geographically clustered, with dense concentrations along the Saudi Red Sea coast and the eastern Mediterranean, whereas arid interior ecosystems and sub-Saharan transitional habitats remain systematically underrepresented (Coleine et al., 2024; Leung et al., 2025).

These patterns are partly a product of methodological choices that shaped data collection across the region. Amplicon-based runs account for 80.6% of the corpus, shotgun metagenomics for 16.9%, and metatranscriptomics for 4.8%, reflecting a legacy of cost and throughput considerations that limits the functional questions the database can address (Knight et al., 2018). While amplicon sequencing allows broad community profiling, it lacks direct functional resolution and is constrained by primer bias. Consequently, antimicrobial resistance gene reservoirs, metabolic pathway distributions, and viral community composition cannot be assessed using amplicon data alone (Jovel et al., 2016; Ranjan et al., 2016). Shotgun metagenomics has expanded among leading contributors since 2018; however, the region continues to lag behind global trends in functional metagenomics, emphasizing the need for wider implementation of shotgun approaches (Bharti & Grimm, 2021; Lloyd-Price et al., 2017).

Technology–ecosystem interaction analysis reveals that long-read platforms are not uniformly distributed across sample types. PacBio SMRT’s over-representation in Environmental samples (z = 4.91) is unsurprising given the platform’s dominance in complex metagenomic work. Soil, sediment, and marine communities present assembly challenges that short reads cannot adequately address. Recovery of near-complete MAGs from these habitats depends on the read lengths that long-read technologies provide (Bickhart et al., 2022; Zhang et al., 2023).

ONT tells a different story. Its concentration in Human samples (z = 6.00) owes more to practical deployment advantages than to raw sequencing performance: the platform travels well, outputs data in real time, and has found genuine utility in point-of-care diagnostics where laboratory infrastructure is scarce (Yek et al., 2022) and full-length 16S rRNA profiling (V1–V9) for species-level resolution in gut, oral, and respiratory microbiomes (Curry et al., 2022). Single-nucleotide accuracy is a prerequisite for AMR gene detection and fine-scale variant calling, and early ONT chemistries did not reliably meet this threshold. R10.4.1 has changed that calculus, producing near-finished assemblies that hold up against PacBio HiFi benchmarks (Sereika et al., 2022). BGI/DNBSEQ platforms show the strongest Human-sample bias in the dataset (z = 11.24), a pattern that extends beyond technical adoption trends. Chinese sequencing partnerships expanded substantially across MENA after 2019, with Saudi Arabia representing the most prominent recipient of this investment (Abdill et al., 2022).

Temporally, long-read penetration rose from 0% before 2018 to 6.2% in 2024 (Mann–Kendall τ = 0.78, p = 0.002), mirroring the global inflection point marked by long-read sequencing being named Nature Methods’ Method of the Year for 2022 (Marx, 2023), with Turkey (9.1%), Saudi Arabia (7.4%), and Egypt (5.8%) leading regional adoption. Nevertheless, adoption across MENA remains well below the penetration rates seen in high-income countries with mature sequencing ecosystems (Agustinho et al., 2024), held back by higher per-sample costs, more stringent DNA quality requirements, a limited supply of validated bioinformatics pipelines, and restricted access to high-performance computing (Agustinho et al., 2024; Yek et al., 2022). Unlocking the full potential of long-read metagenomics in the region will depend on dedicated investment in platform accessibility, bioinformatics training, and the construction of MENA-specific reference databases (Arif et al., 2025; Olsen & Riber, 2025).

The thematic structure of the database provides important context for these patterns. NLP-based K-means clustering (k=9, silhouette=0.590) and cross-tabulation analyses (χ²=35,969.3, p<1e-300) converge on the same conclusion: national identity strongly predicts research thematic focus. Saudi Arabia is statistically over-represented in environmental metagenomics (marine, coral, desert); Turkey in clinical research; and Morocco and Iran in plant-associated and agricultural microbiomes. IndVal analysis independently confirms these signatures through country-specific indicator metagenome types: Lebanon’s food metagenomes (IndVal=0.648), Jordan’s chicken gut (0.596), Bahrain’s sludge (0.571). National research priorities thus reflect local environments alongside funding priorities (Behzad et al., 2016; Currie-Alder et al., 2018).

At the sample level, seawater and marine (n = 9,912), soil and desert (n = 9,441), stool and gut (n = 6,467), oral and saliva (n = 2,310), and wastewater and sludge (n = 2,104) are the five most prevalent categories. Saudi Arabia is dominated by marine and coral tissue samples reflecting its Red Sea research program (Abuzahrah, 2025; Ghobashy et al., 2025). The most balanced human body-site profiles are found in Turkey, Tunisia, and Qatar and the highest plant-compartment diversity across root, rhizosphere, and phyllosphere niches is found in Morocco, Iran, and Sudan (Chukwuneme & Babalola, 2025; Qu et al., 2020). Plant-associated, clinical, and fungal datasets remain underrepresented, in line with the previously identified limitations in PERMANOVA category power. Countries with established thematic niches could help reduce these gaps by contributing more targeted datasets to global resources on fermented foods.(Obafemi et al., 2022; Yasir et al., 2023), pastoralism-associated microbiomes (Mubaraki, 2025), and arid-zone ecology (Coleine et al., 2024; Leung et al., 2025).

Collectively, these results provide a coherent, quantitatively supported overview of MENA metagenomics as a field that is expanding quickly but unevenly. This development appears to reflect a combination of real ecological diversity, distinct national research priorities, and ongoing disparities in research capacity and data governance. The agreement between ecological, spatial, statistical, and text-mining analyses across similar patterns adds robustness to the findings and highlights the usefulness of integrated, database-level approaches for describing regional scientific landscapes.

## 5. LIMITATIONS

Although overall metadata completeness averages 73.97% under a MIxS-MIMS proxy framework, this figure masks severe gaps in analytically essential variables: geographic coordinates (latitude/longitude) are reported in fewer than 15% of runs, environmental descriptors in approximately 25%, host phenotype fields in 10–30%, and DNA extraction method in fewer than 15%, consistent with well-documented patterns of incomplete metadata submission across public repositories (Kim et al., 2025; Vangay et al., 2021; Zass et al., 2024). GPS validation identified 577 runs (1.35% of coordinate-bearing entries) where recorded locations did not correspond to stated country origins, a finding that supports incorporating routine coordinate verification into standard preprocessing workflows ahead of any spatial analysis(Deck et al., 2017).

All ecological and compositional analyses are conducted at the metadata level without access to sequence-level taxonomic or functional profiles; PERMANOVA results therefore reflect variation in sampled ecosystem types rather than direct microbial community composition and should be interpreted as descriptive patterns.

The database includes only publicly available SRA entries and excludes private or unpublished datasets, a constraint that is widely reported in microbiome repositories, where many studies either do not release raw data or provide insufficient metadata for reuse ((Jurburg et al., 2020; Kelliher et al., 2025). The 1,171-day median gap between sample collection and public limits the relevance of these data for outbreak monitoring and time-sensitive epidemiological work, and points to broader structural delays in regional data-sharing systems (Jurburg et al., 2020; Kelliher et al., 2025). Within the MENA context, these delays are compounded by limited institutional capacity for retrospective metadata curation, a problem concentrated among low-volume contributors (Cernava et al., 2022; Zass et al., 2024).

Addressing these issues requires concrete movement toward FAIR data principle compliance (Wilkinson et al., 2016), including standardized journal submission forms, nationally appointed data stewards, and coordinated regional biobanking infrastructure.

## 6. FUTURE DIRECTIONS

The MENA Microbiome Database is intended as a living resource. Planned development includes automated metadata refresh pipelines, enrichment protocols, and integration of shotgun sequence data to support functional analysis at scale. Constructing a dedicated MENA AMR gene catalog represents the most immediate priority, given mounting concern over resistance transmission across environmental, agricultural, and clinical settings. Clinical and fungal microbiomes represent the most acute sampling gaps in the dataset: effect sizes are substantial, yet sample counts remain critically low. Broader geographic coverage presents a different kind of challenge. Libya, Syria, Palestine, Yemen, and South Sudan are all severely undersampled, and closing these gaps will require sustained international funding and cross-institutional collaboration rather than piecemeal national efforts. The database itself was built as a single-file HTML resource to minimize technical barriers to access and reuse across the region.

## 7. CONCLUSION

The MENA Microbiome Database is the first comprehensive, harmonized metagenomic resource compiled for the Middle East and North Africa, and offers a foundational infrastructure for microbiome research across the region. Through integrated ecological, spatial, statistical, and text-mining analyses applied to 60,126 runs from 24 countries, this work delivers an empirical map of how microbiome research is distributed, structured, and constrained across this geographically and ecologically diverse macro-region.

Geography significantly structures environmental, animal-associated, and plant-associated microbiomes but not human-associated microbiomes; national identity strongly predicts research thematic focus; and sampling effort remains spatially clustered around a small number of high-infrastructure hotspots. Geographic coverage, sample type diversity, functional resolution, and metadata quality each represent unresolved deficiencies that should guide the next phase of MENA microbiome research. That multiple independent analytical approaches converge on consistent, interpretable patterns strengthens confidence in these findings and points to the broader utility of integrative, database-level methods for mapping regional research landscapes and addressing persistent inequities in global microbiome science.

## Supporting information

MENA_MICROBIOME_Platform

## Consent for publication

Not applicable.

## Data Availability

All data, code, and the interactive platform are openly available. The GitHub repository (https://github.com/nour0810/mena-microbiome-db) contains all analysis scripts and platform source code. The web platform is hosted at https://nour0810.github.io/mena-microbiome-db/. Raw metagenomic records are available from ENA and NCBI SRA under their original accession numbers.

## Authors’ contributions

N.E.H.M designed the study and conducted the analysis. M.G-B.A, I.B and R.D contributed to data acquisition and analysis. N.E.H.M, M.G-B.A, I.B and R.D prepared and wrote the original draft. L.A.K and R.G supervised the study and revised the manuscript. All authors read and approved the final manuscript.

